# Cross regulation in a three-component cell envelope stress signaling system of *Brucella*

**DOI:** 10.1101/2023.04.15.536747

**Authors:** Xingru Chen, Melene A. Alakavuklar, Aretha Fiebig, Sean Crosson

**Affiliations:** Department of Microbiology and Molecular Genetics, Michigan State University, East Lansing, Michigan USA

## Abstract

A multi-layered structure known as the cell envelope separates the controlled interior of bacterial cells from a fluctuating physical and chemical environment. The transcription of genes that determine cell envelope structure and function is commonly regulated by two-component signaling systems (TCS), comprising a sensor histidine kinase and a cognate response regulator. To identify TCS genes that contribute to cell envelope function in the intracellular mammalian pathogen, *Brucella ovis*, we subjected a collection of non-essential TCS deletion mutants to compounds that disrupt cell membranes and the peptidoglycan cell wall. Our screen led to the discovery of three TCS proteins that coordinately function to confer resistance to cell envelope stressors and to support *B. ovis* replication in the intracellular niche. This tripartite regulatory system includes the known cell envelope regulator, CenR, and a previously uncharacterized TCS, EssR-EssS, which is widely conserved in *Alphaproteobacteria*. The CenR and EssR response regulators bind a shared set of sites on the *B. ovis* chromosomes to control transcription of an overlapping set of genes with cell envelope functions. CenR directly interacts with EssR and functions to stimulate phosphoryl transfer from the EssS kinase to EssR, while CenR and EssR control the cellular levels of each other via a post-transcriptional mechanism. Our data provide evidence for a new mode of TCS cross-regulation in which a non-cognate response regulator affects both the activity and protein levels of a cognate TCS protein pair.

**Importance:** As intracellular pathogens, *Brucella* must contend with a variety of host-derived stressors when infecting a host cell. The inner membrane, cell wall, and outer membrane — i.e. the cell envelope — of *Brucella* provides a critical barrier to host assault. A conserved regulatory mechanism known as two-component signaling (TCS) commonly controls transcription of genes that determine the structure and biochemical composition of the cell envelope during stress. We report the identification of previously uncharacterized TCS genes that determine *B. ovis* fitness in the presence of cell envelope disruptors and within infected mammalian host cells. Our study reveals a new molecular mechanism of TCS-dependent gene regulation, and thereby advances fundamental understanding of transcriptional regulatory processes in bacteria.

## Introduction

Genes that regulate adaptation to changes in the physical and chemical state of the environment are important fitness determinants in bacteria (1-6). As intracellular pathogens, *Brucella* species inhabit the interior of mammalian cells and must adapt to numerous host-derived assaults including oxidative stress, nutrient limitation, and low pH. Accordingly, *Brucella* strains harboring mutations in genes that function to mitigate these stresses often have reduced fitness in the intracellular niche, e.g. (7-17). As in other bacteria, two-component signaling systems (TCS) are prominent regulators of *Brucella* stress responses (18, 19). Canonical two-component signaling entails signal-induced autophosphorylation of a sensor histidine kinase (HK), followed by specific phosphoryl transfer to a single DNA-binding response regulator (RR) protein that controls a transcriptional response (20). The vast majority of systems follow this classic paradigm, though there are examples of signal transduction processes that likely involve cross-regulation between otherwise distinct TCS protein pairs, e.g. (21-27).

A common function of TCS pathways is to regulate the biochemical composition and transport capacity of the cell envelope (28, 29), which serves as the barrier between the interior of the cell and the surrounding environment (30). As Gram-negative bacteria, the envelope of *Brucella* spp. contains two structurally distinct lipid bilayers with a peptidoglycan cell wall in the thin periplasmic space between. Only two of the six classical *Brucella* species, *B. canis* and *B. ovis*, are naturally “rough”, meaning they do not synthesize repeating O-polysaccharide units that extend from the core LPS oligosaccharide of the outer membrane (31). These rough species are not as commonly studied as “smooth” *Brucella* species, which synthesize O-polysaccharide and are the major zoonotic pathogens. *B. ovis* therefore provides an interesting comparative model to investigate genes that regulate molecular features of the *Brucella* cell envelope. We sought to define the contribution of non-essential *B. ovis* TCS genes to cell survival under cell envelope stress conditions, with an additional goal of assessing functional relationships between TCS genes. To this end, we generated a collection of mutant strains harboring in-frame deletions of 32 non-essential TCS genes and measured growth and viability phenotypes of these strains on solid media containing sodium dodecyl sulfate (SDS) or carbenicillin, to target the cell membrane and cell wall, respectively. Five *B. ovis* TCS mutant strains were either more sensitive or more resistant than wild type (WT) to these specific cell envelope disruptors. Among these mutants, strains lacking the RR encoded by locus *BOV_1929* (BOV_RS09460; *cenR*) or the RR and HK encoded by locus *BOV_1472*-*73* (BOV_RS07250-55; *essR-essS*) had similar phenotypes under the tested conditions. Through a series of genetic, genomic, and biochemical experiments, we uncovered evidence for direct cross-regulation between these three TCS proteins, which control transcription of a common gene set to support *B. ovis* replication under envelope stress conditions and in the host intracellular niche. Specifically, the conserved alphaproteobacterial cell envelope regulator, CenR (32-34), stimulates phosphoryl transfer between the cognate RR-HK pair, EssR-EssS, while CenR and EssR regulate the steady-state levels of each other in *B. ovis*. Our results expand understanding of molecular mechanisms of two-component signal transduction and define an unusual mode of TCS cross-regulation in *Brucella* that controls gene expression to support envelope stress resistance and intracellular replication.

## Results

### Tn-seq identifies a set of candidate essential TCS genes in Brucella ovis

There are 47 TCS genes in the *B. ovis* genome (Genbank accessions NC_009504 and NC_009505); four of these genes contain frameshift mutations and are defined as pseudogenes (Table S2). Several histidine kinases and response regulators are reported to be essential in *Brucella* spp. based on previously published Tn-seq data in *B. abortus* (35) and we sought to comprehensively define the essential set of TCS genes in *B. ovis* using *himar* transposon sequencing. Analysis of Tn-*himar* insertion sites in a collection of approximately 200,000 *B. ovis* mutants revealed » 54,000 unique TA dinucleotide insertions in the *B. ovis* genome (36). We used Hidden Markov Model and Bayesian-based approaches (37) to identify candidate essential genes based on this Tn-seq dataset (Table S1). As expected, many of the known TCS regulators of *Brucella* cell cycle and cell development (38) are defined as essential using this approach including *ctrA, cckA, chpT, pdhS, divL*, and *divK*. Though insertion counts were lower in the developmental/cell cycle regulator *cpdR* relative to local background, this gene did not reach the essential threshold by our analysis. The *bvrS*-*bvrR* two-component system, which is homologous to the *chvG*-*chvI* system of other Alphaproteobacteria (39), is also essential. This result is consistent with a report by Martín-Martín and colleagues that the *bvrSR* genes cannot not be deleted in *B. ovis* (40). Notably, the *bvrSR* system is not essential in the smooth strains, *B. melitensis* (39) and *B. abortus* (41). Finally, the *ntrX* regulator is also essential. Considering a recent report of strong genetic interactions between *ntrX, chvG*, and *chvI* in *Caulobacter* (42), the essential phenotypes of *bvrS-bvrR* and *ntrX* may be related.

As this study is focused on cell envelope regulation in a rough *Brucella* species, we note that several genes with cell envelope functions were identified as essential in *B. ovis* that are not essential in the smooth species, *B. abortus* (35), including a putative L,D-transpeptidase (*BOV_0757*) and a putative D-alanyl D-alanine carboxypeptidase (*BOV_1129*). Additionally, a DegQ-family serine endoprotease (*BOV_0610*), hydroxymethylpyrimidine phosphate kinase ThiD (*BOV_0209*), the putative magnesium transporter MgtE (*BOV_A0822*), the iron sulfur cluster insertion protein ErpA (*BOV_0886*), and the RNA chaperone Hfq (*BOV_1070*) are essential in *B. ovis* but not in *B. abortus* (35).

### TCS mutants sensitive (and resistant) to cell envelope disruptors in vitro

We next generated a set of 24 mutant strains harboring in-frame deletions of 32 non-essential TCS genes in *B. ovis* (Table S2), comprising 8 double mutants and 16 single mutants. In cases where sensor kinase and response regulator genes are adjacent in the *B. ovis* genome, we inferred that these genes functioned together and therefore deleted both. The double deletion strains include: *ΔBOV_0603-0604* (*feuPQ)*, Δ*BOV_0611-0612*, Δ*BOV_1472-1473, ΔBOV_1075-1076* (*ntrCB*), *ΔBOV_A0037-A0038* (*nodVW*), *ΔBOV_A209-A210, ΔBOV_0357-0358*, and *ΔBOV_A0412-A0413*. The single deletion strains include: *ΔBOV_0099, ΔBOV_0131 (regA), ΔBOV_0289, ΔBOV_0312 (phoR), ΔBOV_0331 (prlR), ΔBOV_0332 (prlS), ΔBOV_0577 (divJ), ΔBOV_0615 (pleC), ΔBOV_1602, ΔBOV_1604 (phyR), ΔBOV_1929 (cenR), ΔBOV_2058 (phoB), ΔBOV_A0358, ΔBOV_A0554 (lovhK), ΔBOV_A1045 (ftcR), ΔBOV_A0575 (pleD)*, (Table S2). To identify TCS genes that function to regulate the cell envelope, we subjected this collection of non-essential deletion mutants to treatments known to compromise envelope integrity. Specifically, WT and *B. ovis* mutant strains were serially diluted and plated on plain tryptic soy agar supplemented with 5% sheep blood (TSAB) and on TSAB containing *a)* the detergent sodium dodecyl sulfate (SDS) or *b)* the cell wall targeting (beta-lactam) antibiotic carbenicillin. Five of the 24 mutant strains showed a growth or viability phenotype that differed from WT in multiple conditions including Δ*BOV_0577*, Δ*BOV_1602*, Δ*BOV_1929*, Δ*BOV_0611-0612*, and Δ*BOV_1472-1473* (Figures 1 and S1).

**Figure 1:**
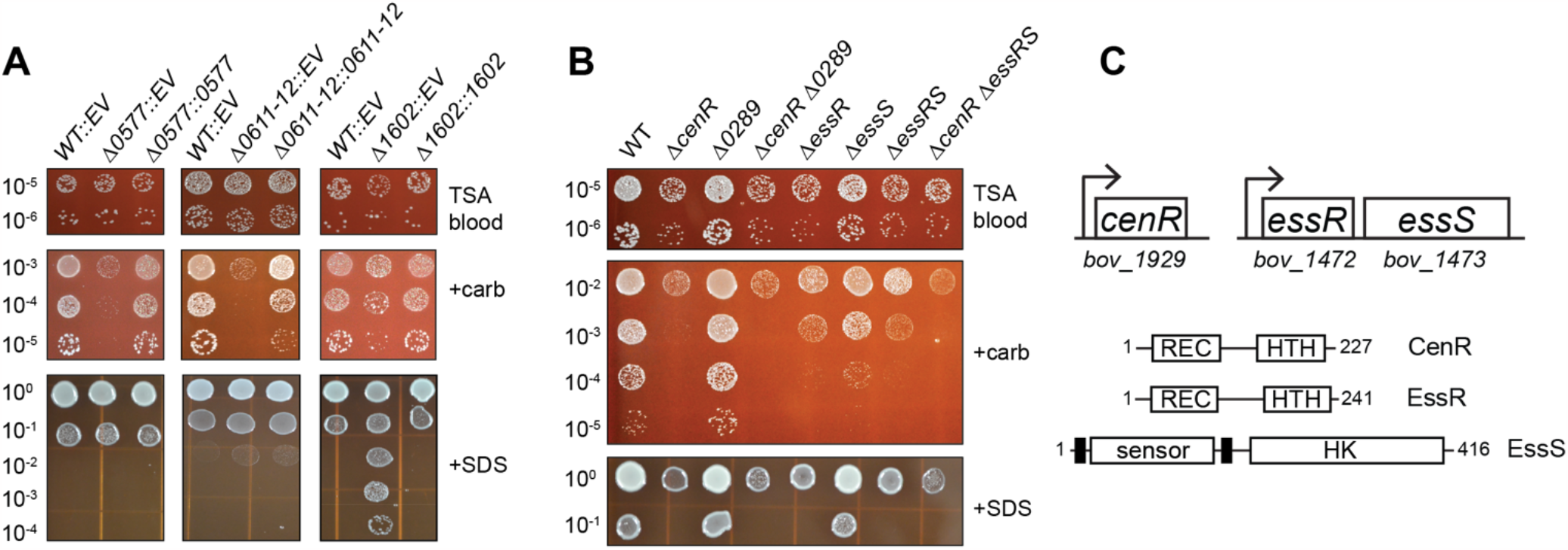
Analysis of *B. ovis* TCS deletion mutants identified five genes/operons that contribute to carbenicillin and SDS resistance *in vitro*. (A) Strains harboring in-frame unmarked deletions (D) of *B. ovis* TCS genes *BOV_0577, BOV_0611-0612*, and *BOV_1602*, carrying integrated empty vectors (EV) or genetic complementation vectors (::*gene locus number*) were plated in log_10_ dilution series on plain TSA blood agar (TSAB), TSAB containing 2 µg/ml carbenicillin (+carb) or TSAB containing 0.0045% SDS (+SDS). (B) Strains harboring in-frame unmarked deletions (D) of *B. ovis* TCS gene loci *BOV_1929* (*cenR*), *BOV_0289, BOV_1472* (*essR*), *BOV_1473* (*essS*), alone and in combination plated as in panel A. Genetic complementation of panel B strains using a lower SDS concentration is shown in Figure S1. Dilution plating experiments were repeated at least three times for all strains, and one representative experiment is shown. (C) (top) Cartoon of the *cenR* and *essR-essS* genetic loci with *bov* gene locus numbers. (bottom) Protein domain models of CenR, EssR, and EssS with number of amino acid residues in each protein. Domains are labeled as follows: REC - receiver domain; HTH - helix-turn-helix DNA binding domain, sensor - extracellular sensor domain, HK - histidine kinase domain. Predicted transmembrane helices flanking the sensor domain are indicated by heavy black lines.

All five mutants with envelope stress defects had reduced viability on TSAB containing 2 mg/ml carbenicillin relative to WT (Figure 1 and Figure S1), though the carbenicillin sensitivity of Δ*BOV_1602* was less severe than other strains (Figure 1A). The Δ*BOV_1929* single mutant and Δ*BOV_1472-1473* double mutant had growth defects on plain TSAB, as evidenced by smaller colony size (Figure 1B and Figure S1B). Growth of WT *B. ovis* was impaired on TSAB containing 0.0045% SDS, but Δ*BOV_1929* and Δ*BOV_1472-1473* were even more severely impacted under this condition (Figure 1B). Conversely, deletion of *BOV_1602*, a cytoplasmic HWE-family histidine kinase (43) containing an amino-terminal PAS domain (44), strongly enhanced *B. ovis* viability on SDS plates: growth was apparent at titers approximately three log_10_ units more diluted than WT (Figure 1A). *BOV_1602* is adjacent to the *nepR*-*ecfG*-*phyR* general stress response (GSR) locus (8) in *B. ovis*. It was reported that orthologs of *BOV_1602* do not regulate *Brucella* spp. GSR *in vitro* or *in vivo* (7, 45), but control of GSR transcription is complex (2, 46) and may involve *BOV_1602* under select conditions. In all cases, the observed agar plate phenotypes were genetically complemented by integration of a single copy of the deleted gene(s) with their native promoter at the *glmS* locus (Figure 1 and Figure S1A).

### A genetic connection between BOV_1472-1473 (essRS) and BOV_1929 (cenR)

There was notable congruence in the agar plate growth defects of the Δ*BOV_1929* and Δ*BOV_1472-1473* strains across the tested conditions, though Δ*BOV_1929* was more sensitive to carbenicillin than Δ*BOV_1472-1473* (Figure 1B). BOV_1929 has high sequence identity to CenR, which is a known regulator of cell envelope structure in the Alphaproteobacteria (33, 34): *BOV_1929* and *Caulobacter crescentus, Rhodobacter sphaeroides*, and *Sinorhizobium meliloti* CenR are reciprocal top-hit BLAST pairs (ranging from 65-80% identity over the full length of the protein). We have thus named *BOV_1929, cenR*. Consistent with our data, *Brucella melitensis cenR* (also known as *otpR*) has a reported role in beta-lactam tolerance (47) and acid stress response (48). Studies in *C. crescentus* and *R. sphaeroides* have identified the histidine kinase CenK as the cognate regulator of CenR (33, 34), but the *B. ovis* genome does not encode a protein with high sequence identity to *Caulobacter* or *Rhodobacter* CenK. This is consistent with a previous report that *B. melitensis* does not encode an evident CenK ortholog (49). The *B. ovis* sensor HK most closely related to *C. crescentus* and *R. sphaeroides* CenK is encoded by gene locus *BOV_0289* (33-37% amino acid identity). *BOV_0289* encodes the most likely HK partner for CenR based on a Bayesian algorithm to predict TCS HK-RR pairs (50). However, the *ΔBOV_0289* deletion mutant grew the same as WT in the tested stress conditions and a *ΔBOV_0289 ΔcenR* double mutant phenocopied the *ΔcenR* single mutant (Figure 1B). These data indicate that BOV_0289 is not an activator of CenR in these conditions. Given the **e**nvelope **s**tress **s**urvival phenotypes of the Δ*BOV_1472-1473* double deletion mutant, we hereafter refer to the DNA-binding response regulator gene *BOV_1472* as *essR* and the sensor histidine kinase gene *BOV_1473* as *essS*. To our knowledge, EssS and EssR have not been functionally characterized, though EssS is 68% identical to the RL3453 sensor kinase that has been functionally linked to *Rhizobium leguminosarum* plant root attachment (51). EssR is an OmpR-family response regulator, while the EssS sensor kinase has two predicted transmembrane helices and primary structure features that resemble the well-studied CpxA and EnvZ cell envelope regulators (52-54). To assess phenotypic relationships between the Δ*essR-essS* and Δ*cenR* mutants, we generated *B. ovis* strains harboring in-frame deletions of either *essR* or *essS* and a *ΔcenR ΔessR-essS* triple mutant and evaluated growth of these strains on TSAB containing SDS or carbenicillin as above. The *ΔessR* strain phenocopied the *ΔessR-essS* double mutant in all conditions. The *ΔessS* mutant was indistinguishable from WT in the presence of 0.0045% SDS, however, at a lower SDS concentration (0.004%) *ΔessS* showed an intermediate sensitivity phenotype (Figure S1B). *ΔessS* also showed an intermediate carbenicillin sensitivity phenotype. The *ΔcenR ΔessR-essS* triple mutant phenocopied the *ΔcenR* and *ΔessR* single mutants (Figure 1B & S1B). We further evaluated *ΔcenR, ΔessR*, and *ΔessS* mutants in three additional stress conditions that perturb the cell envelope in different ways: ethylenediaminetetraacetic acid (EDTA) is a divalent cation chelator that can destabilize the outer membrane (55); polymyxin B is a cationic antimicrobial peptide that targets the outer membrane; and NaCl can function as an osmotic stressor that disrupts envelope integrity. *ΔcenR* and *ΔessR* again phenocopied each other in all three treatment conditions. Both response regulator mutants were more sensitive to EDTA and polymyxin B, and both were more resistant to NaCl compared to wild type. These results provide further support that CenR and EssR function in the same pathway. Like the RR mutants, *ΔessS* was sensitive to EDTA and polymyxin B. However, *ΔessS* was not NaCl resistant (Figure S2). From these results, we hypothesize that the EssS sensor kinase has distinct regulatory roles in different stress conditions. The sequence relatedness between EssR and CenR is moderate when compared to pairings of all *B. ovis* response regulators with DNA binding domains, with 32% identity and 50% similarity. Specific regions of identity and similarity in their primary structures are presented in Figure S1C.

### Contribution of the CenR and EssR aspartyl phosphorylation sites to stress survival

CenR and EssR are both DNA-binding response regulator proteins, which are typically regulated by phosphorylation of a conserved aspartic acid residue in the receiver domain. To assess the functional role of the CenR aspartyl phosphorylation site (D55), we tested whether expression of non-phosphorylatable (*cenR*_*D55A*_) or putative phosphomimetic (*cenR*_*D55E*_) alleles of *cenR* could genetically complement the defects of the *ΔcenR* strain in the presence of SDS or carbenicillin. Both mutant alleles of *cenR* restored *ΔcenR* growth/survival to WT levels, indicating that the CenR phosphorylation site does not impact CenR function under these conditions (Figure 2). We conducted the same experiment for EssR, testing whether *essR*_*D64E*_ and *essR*_*D64A*_ could genetically complement the *ΔessR* growth defects. *essR*_*D64E*_ expression restored wild-type like growth to *ΔessR* on plain TSAB and TSAB-SDS plates but resulted in a strain that was even more sensitive than *ΔessR* to carbenicillin. Expression of *essR*_*D64A*_ failed to complement *ΔessR* under all conditions (Figure 2). We conclude that EssR phosphorylation at D64 contributes to *in vitro* stress survival of *B. ovis*.

**Figure 2:**
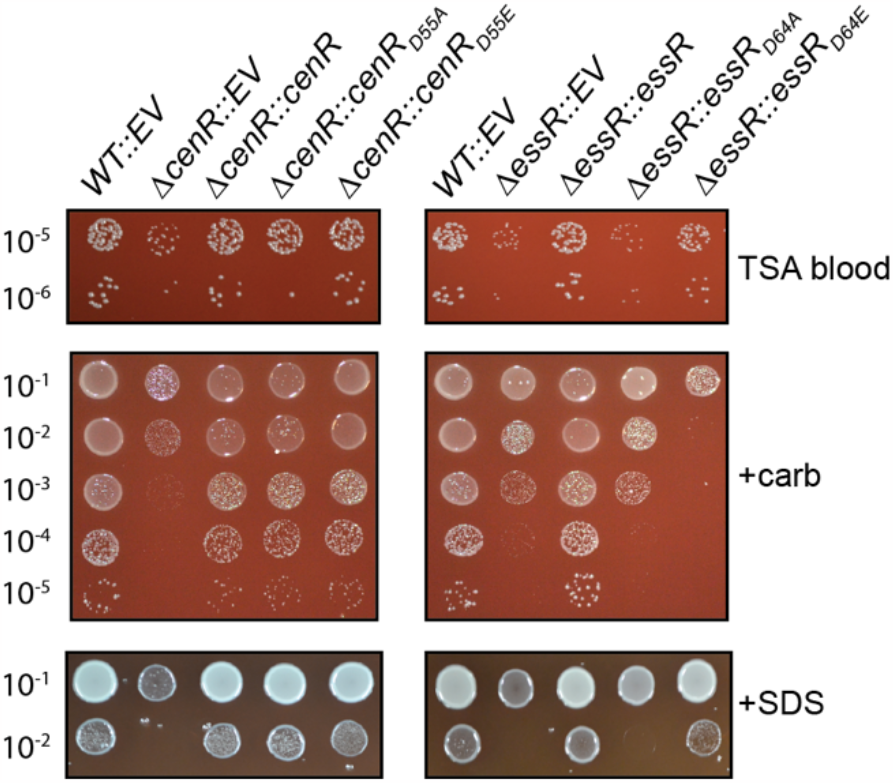
Functional analysis of the conserved aspartyl phosphorylation sites of the CenR and EssR response regulators. Test for genetic complementation of D*cenR* and D*essR* envelope stress phenotypes by expression of either non-phosphorylatable alleles of *cenR* (D55A) and *essR* (D64A) or putative phosphomimetic alleles (D55E / D64E). Conditions and dilution plating as in Figure 1; Plain TSA blood (TSAB) plates, TSAB + 2 µg/ml carbenicillin (+carb) and TSAB + 0.0045% SDS (+SDS). D*cenR* and D*essR* each carry an empty vector as control (EV). Experiments were repeated at least three times for all strains and one representative experiment is shown.

### cenR *and* essR *mutants have equivalent cell size defects*

Previous studies of CenR in *C. crescentus* and *R. sphaeroides* have shown that depletion or overexpression of *cenR* leads to large defects in cell envelope structure and/or cell division (33, 34). Considering these results and the sensitivity of *B. ovis ΔcenR* to SDS and carbenicillin, we inspected *ΔcenR, ΔessR*, and *ΔessS* cells by phase contrast light microscopy and cyro-electron microscopy for defects in cell envelope structure or cell morphology. Deletion of *cenR, essR* or *essS* did not result in apparent changes in cell morphology or cell division as assessed by phase contrast microscopy at 630x magnification (Figure S3A). However, an analysis of cell size revealed that both *ΔcenR* and *ΔessR* mutant cells were larger than WT; the average area of the mutant cells was approximately 12% greater than WT in 2D micrographs (p<0.0001) (Figure S3B). Again, the parallel phenotype of *ΔcenR* and *ΔessR* supports a model in which these two genes execute related functions. An intact phosphorylation site was not required for CenR or EssR to affect cell size, as expression of either phosphorylation site mutant allele of *cenR*(D55) or *essR*(D64) restored cell size to WT levels (Figure S3B). We did not observe major cell membrane defects in *ΔcenR* and *ΔessR* mutant cells by cryo-EM (Figure S3C). We conclude that the SDS and carbenicillin resistance defects we observe in the D*cenR* and D*essR* strains are associated with a defect in cell size control.

### *B. ovis ΔcenR, ΔessR*, and *ΔessS* strains have equivalent fitness defects in a macrophage infection model

As *B. ovis* is an intracellular pathogen, we sought to examine the importance of *cenR, essR* and *essS* in the intracellular niche. The interior of mammalian phagocytes more closely models conditions encountered by the bacterium in its natural host, and many mutants with defects in cell envelope processes are attenuated in infection models (56). *ΔcenR, ΔessR*, and *ΔessS* strains had no defect after entry (2h post-infection (p.i.)) or at early stages of infection (6h & 24h p.i.) of THP-1 macrophage-like cells relative to WT. By 48h p.i., recoverable CFU of WT *B. ovis* increased, consistent with adaptation and replication in the intracellular niche; CFU of *ΔcenR, ΔessR*, and *ΔessS* did not increase appreciably between 24 and 48h. The 1 log_10_ unit intracellular replication defect of the *ΔcenR, ΔessR*, and *ΔessS* strains at 48h was complemented by expression of the deleted gene(s) from an ectopic locus (Figure 3). Thus, we conclude that *cenR, essR*, or *essS* do not affect macrophage entry or early survival but all three genes similarly affect replication and/or survival after 24h. These results indicate that *essS, essR*, and *cenR* contribute to *B. ovis* fitness after the establishment of the replicative niche inside the *Brucella*-containing vacuole.

**Figure 3:**
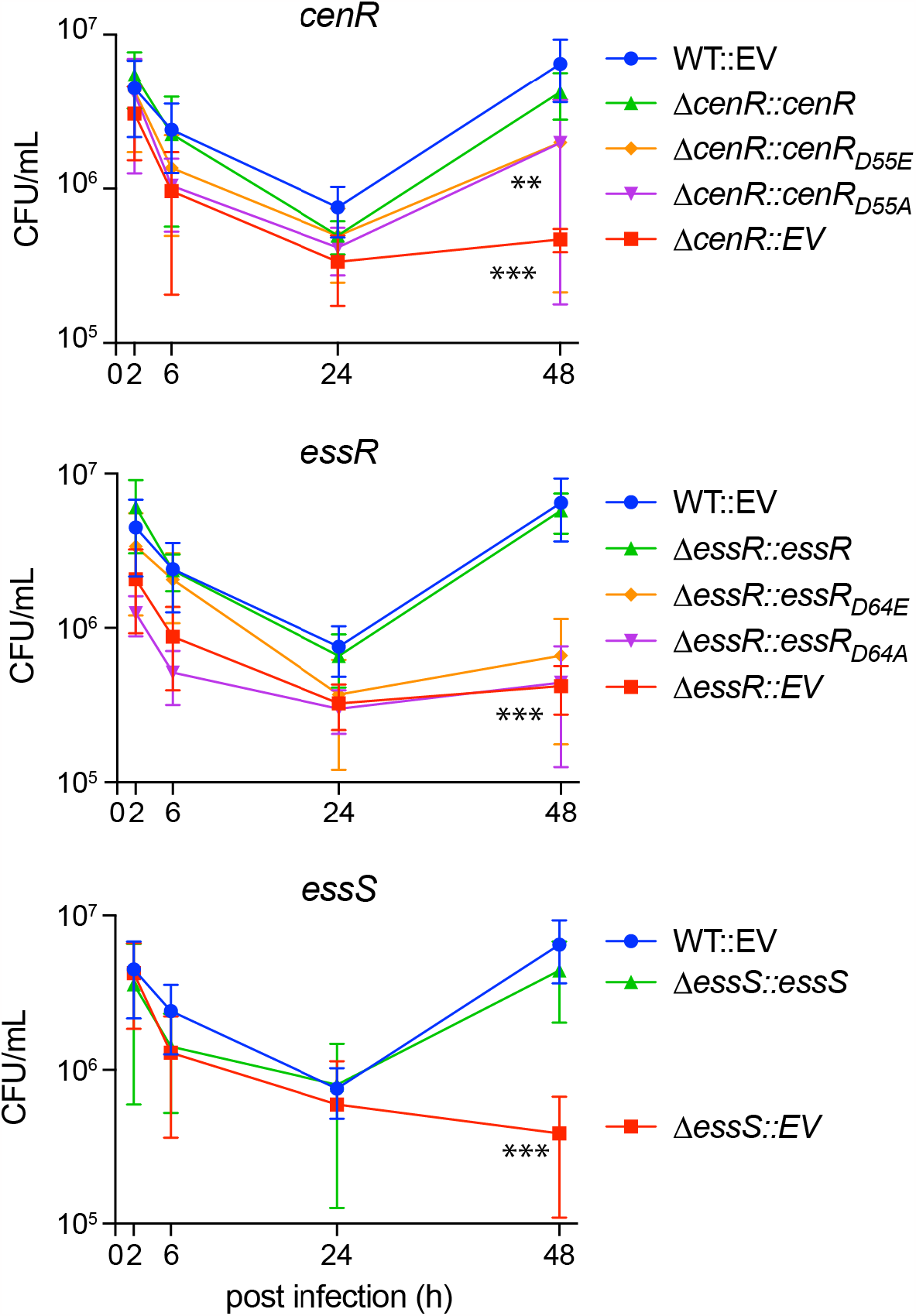
*B. ovis ΔcenR*, Δ*essR*, and Δ*essS* deletion strains have reduced fitness in the intracellular niche of mammalian macrophage-like cells; intact CenR and EssR aspartyl phosphorylation sites contribute to *B. ovis* fitness in the intracellular niche. Log_10_ colony forming units (CFU) per well of wild-type *B. ovis* carrying an integrated empty vector (WT::EV) (blue), Δ*cenR*, Δ*essR*, and Δ*essS* carrying an EV (red), or Δ*cenR*, Δ*essR*, and Δ*essS* expressing the missing gene from an integrated vector (green). Brucellae were isolated from infected THP-1 cells and enumerated at 2, 6, 24 and 48 hours (h) post infection. The contribution of the conserved aspartyl phosphorylation sites of CenR and EssR to intracellular fitness was assessed by testing for genetic complementation of D*cenR* and D*essR* phenotypes by integration of either non-phosphorylatable alleles of *cenR* (D55A) and *essR* (D64A) (purple) or putative phosphomi-metic alleles (D55E / D64E) (orange). Infections were repeated 3 times; error bars represent standard deviation of the three biological replicates. Statistical significance was calculated at 48h p.i. using one-way ANOVA, followed by Dun-nett’s multiple comparisons test to WT::EV control (p < 0.001, **; p < 0.0001, ***).

Additionally, we tested the infection phenotypes of strains harboring alleles of *essR* and *cenR* in which the conserved aspartyl phosphorylation site was mutated. Expression of *cenR*_*D55A*_ or *cenR*_*D55E*_ partially - and equivalently - complemented the 48-hour infection defect of *ΔcenR* (Figure 3A). Expression of either *essR*_*D64E*_ or *essR*_D64A_ failed to complement *ΔessR* in this assay (Figure 3B). We conclude that an intact aspartyl phosphorylation site in both the CenR and EssR receiver domains is required for WT levels of replication in a THP-1 macrophage infection model.

### *cenR, essS* and *essR* mutants are not sensitive to low pH

The Brucella containing vacuole (BCV) is acidified at early time points after infection (1 to 10 hours) in J774 murine macrophages and HeLa cells (57, 58). We observed no defect in recoverable CFU of our mutants at the 2h time point in THP-1 macrophage-like cells (Figure 3), which suggested that acid tolerance was not perturbed in *ΔcenR, ΔessR*, or *ΔessS*. Nonetheless, we sought to more rigorously test whether sustained exposure to acid impacted the survival of *ΔcenR, ΔessR*, and *ΔessS* strains. The acidified BCV has a pH in the 4.0-4.5 range (59), so we tested whether exposure to acidified Brucella broth (pH 4.2) for 2 hours differentially impacted mutant viability. We did not observe significant differences in viability between WT and mutant strains after *in vitro* acid exposure (Figure S4) and conclude that sensitivity to acid in the BCV cannot alone explain the intracellular defects of *cenR, essS* and *essR* mutants. The slower growth rates of *cenR* and *essRS* mutants, which are evident on solid media (Figures 1 & 2 and S1) and in broth (k =0.0028 min^-1^ for WT, 0.0021 min^-1^ for Δ*cenR*, and 0.0022 min^-1^ for Δ*essRS*), could be the primary determinant of their replication defect after 24h. Nonetheless, the more severe defect of the Δ*essS* strain between 24 and 48 hours relative to the other TCS mutants provides evidence that intracellular defects of these TCS mutants are likely complex and multifactorial.

### EssR and CenR regulate a common gene set

Considering that TCS proteins typically function to regulate transcription, we used RNA sequencing (RNA-seq) to assess the relationship between EssRS- and CenR-regulated gene sets. The global transcriptional profiles of *ΔcenR* and *ΔessRS* mutant strains were highly correlated (Figure 4). Filtering genes based on a minimum fold change ^3^ |2| (FDR p-value < 10^−4^) revealed 46 transcription units, containing 53 genes, that were regulated by *essR-essS*. Fifty-two transcription units, containing 61 genes, were regulated by *cenR* (Figure 4; Table S3). These gene sets largely overlap: thirty-eight (38) are differentially expressed in the same direction in both datasets. With few exceptions, genes that met the criteria for differential regulation in only one strain showed similar, but more modest, changes compared to WT in the other strain. These results indicate a high degree of functional overlap of EssRS and CenR, with respect to gene expression. One potential explanation for the observed regulatory overlap is that EssR and CenR transcription depend on each other. However, transcript levels of *essR*-*essS* in a *ΔcenR* background, and *cenR* in *ΔessR-essS* background do not differ significantly from WT (Table S3). We conclude that neither response regulator significantly affects the transcription of the other. Rather, CenR and EssR either independently or coordinately regulate transcription of an overlapping set of *B. ovis* genes.

**Figure 4:**
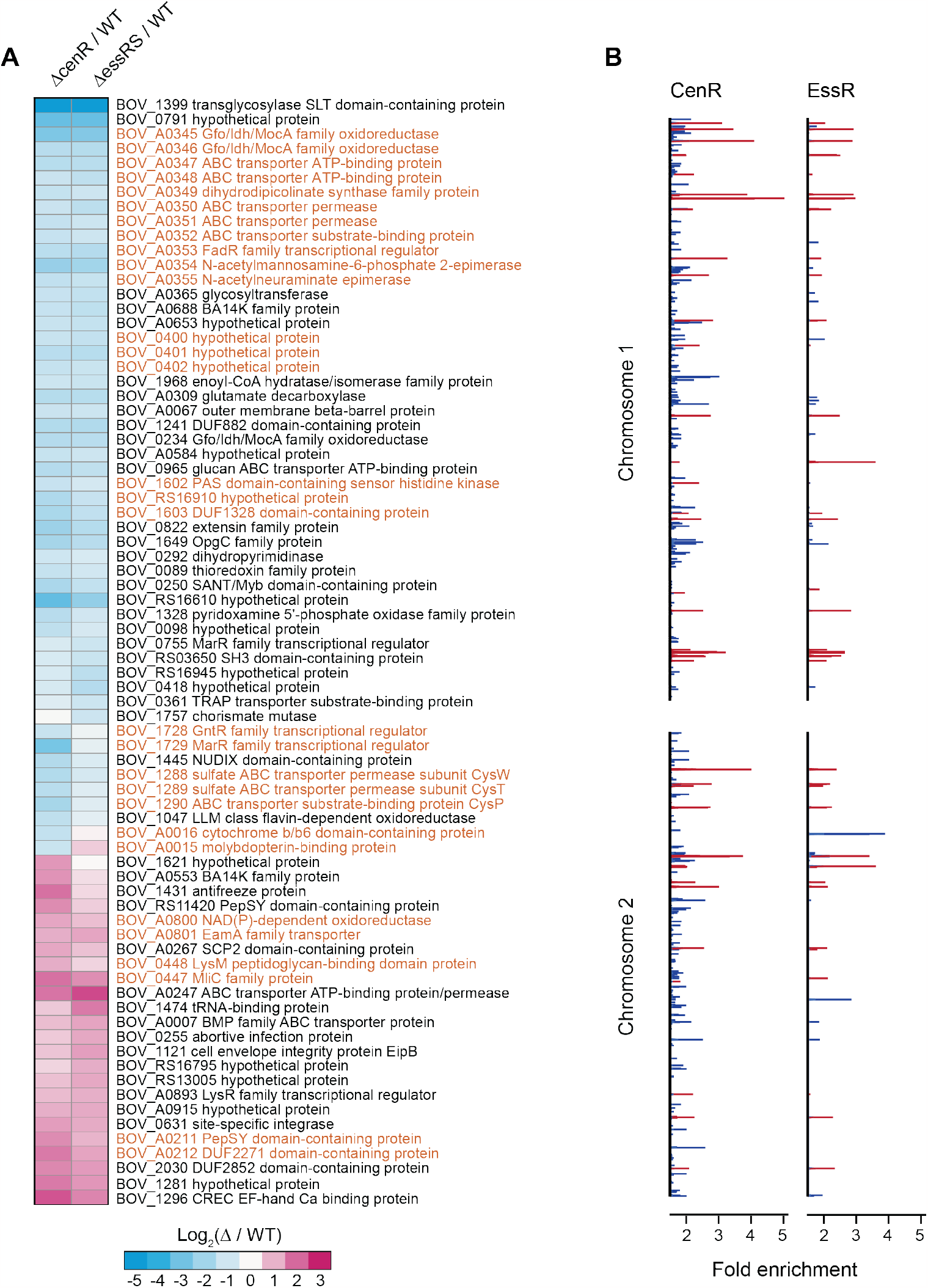
CenR and EssRS regulate an overlapping set of genes and have a correlated ChIP-seq profile. (A) Heat map of log_2_ fold change in gene expression in Δ*cenR* and Δ*essRS* deletion strains relative to wildtype (WT). Genes presented have a fold change ^3^ |2| with an FDR p-value < 10^−4^. Genes highlighted in orange are adjacent on the chromosome. (B) EssR and CenR chromatin immunoprecipitation (ChIP)-seq peaks (q-value £ 0.05; minimum AUC = 20) along chromosomes 1 and 2. Red lines mark significant DNA peaks that are immunoprecipitated by CenR and EssR. Blue lines indicate peaks that are unique to either CenR or EssR.

To further investigate the connection between CenR- and EssR-dependent transcription, we performed chromatin-immunoprecipitation (ChIP)-seq with both EssR and CenR. To promote RR binding to chromosomal target sites, we expressed putative phosphomimetic alleles (DàE) of each RR from their native promoters integrated at the *glmS* locus in each mutant strain; genes were fused to a C-terminal 3xFLAG tag. ChIP of EssR_D64E_-3xFLAG yielded 65 significant peaks across two biological replicates, and ChIP of CenR_D55E_-3xFLAG yielded 47 significant peaks across three biological replicates. Thirty-three peaks were shared between CenR_D55E_ and EssR_D64E_ (Figure 4B and Table S4) providing additional support for a model in which CenR and EssR regulate transcription of a shared gene set, functioning either separately or as a heteromeric complex. We did not observe CenR binding to the *essRS* promoter or EssR binding to the *cenR* promoter in the ChIP-seq data. This is further indication that CenR and EssR do not regulate each other transcriptionally, consistent with the RNA-seq results.

### Genes with cell envelope functions are abundant in the CenR-EssRS regulon

The CenR/EssR regulon prominently features genes encoding membrane transport proteins, several of which are contained in co-regulated clusters (Figure 4). The largest of these clusters encodes carbohydrate metabolism enzymes (BOV_A0354-A0355), transporter sub-units (BOV_A0347-A0348; BOV_A0350-A0352), a DapA-family dihydrodipicolinate synthase/N-acetylneuraminate lyase (BOV_A0349), Gfo/Idh/MocA family oxidoreductases (BOV_A0345-A0346), and a FadR-family transcription regulator (BOV_A0353). Transcription of this entire cluster is significantly reduced following *cenR* or *essRS* deletion (Figure 4; Table S3). Expression of *B. melitensis* orthologs of these genes is highly upregulated in supramammary lymph nodes of goats, specifically at late stages of infection (60) suggesting a role in long-term colonization of animal hosts. Homologs of these genes are also regulated by CenR in *R. sphaeroides* (33) (Table S3), providing evidence for conservation of CenR-dependent transcription across Alphaproteobacterial genera.

Other genes with reduced expression upon *cenR* or *essRS* deletion include a third Gfo/Idh/MocA family oxidoreductase (*BOV_0234*) and the secreted BA14K protein (*BOV_A0688*), which is required for normal spleen colonization in a mouse model of *B. abortus* infection (8). Transcripts of three genes immediately adjacent to the general stress response regulator, PhyR, also decreased upon *cenR* or *essRS* deletion including the previously discussed HWE-family sensor kinase, *BOV_1602*, which impacts *B. ovis* SDS resistance (Figure 1). Additionally, the predicted sulfate ABC transport operon, *cysWTP*, decreased significantly in both deletion mutant strains (Figure 4A; Table S3). Nine genes had significantly higher transcription in *ΔcenR* and *ΔessRS* (Figure 4A; Table S3) including a site-specific DNA integrase (*BOV_0631*), an ABC transporter (*BOV_A0247*), a C-type lysozyme inhibitor (*BOV_0447*) and LysM-domain protein (*BOV_0448*), envelope integrity protein B (*eipB*; *BOV_1121*), and *BOV_1296*, which is directly regulated by the virulence regulator VjbR under select conditions (61). *BOV_1296* encodes acid shock protein 24 (Asp24), which contributes to virulence in later stages of infection in *B. abortus* and *B. melitensis* (62, 63).

*BOV_1399*, encoding a periplasmic soluble lytic murein transglycosylase (SLT) enzyme, exhibited the most significant expression difference in our RNA-seq datasets. The promoter of *BOV_1339* is directly bound by CenR and EssR (Table S4) and transcript levels were approximately 48 times lower than WT in both *ΔcenR* or *ΔessRS* RNA-seq datasets (Figure 4; Table S3). To test whether loss of this predicted cell wall metabolism enzyme contributed to the stress survival phenotypes of *ΔcenR* and *ΔessRS*, we deleted *BOV_1399* and subjected this strain to the same agar plate growth assays described above. However, the phenotypes of *ΔBOV_1399* in these assays were indistinguishable from WT. Notably, the promoters of several genes with known cell envelope functions are bound by CenR, EssR, or both but do not exhibit differential transcription in *ΔcenR* or *ΔessRS*. For example, EssR binds to the promoter of *BOV_0115* (Omp25d), which contributes to cell envelope integrity (64, 65), and *B. ovis*-host interaction (66). The outer membrane autotransporters, *bmaA* and *bmaC*, which are important for protein translocation to the cell surface (67) and for host cell adherence (11) are also bound by CenR/EssR but do not change in expression upon *cenR* or *essRS* deletion. Additional studies are necessary to determine what CenR/EssR-bound or regulated genes determine *B. ovis* cell size and support *B. ovis* fitness under envelope stress and in the intracellular niche.

### CenR and EssR physically interact via their REC domains in a heterologous system

Genetic, transcriptomic, and ChIP-seq data all indicate that the response regulators EssR and CenR directly control expression of a common gene set in *B. ovis* to enable growth in the presence of SDS and carbenicillin. While the molecular processes that determine signaling via a particular TCS pathway are typically insulated from other TCS proteins (20), we postulated that the congruent genetic and molecular phenotypes of *ΔcenR* and *ΔessR* strains could arise through direct molecular interactions of CenR with either EssS, EssR, or both. In fact, in a systematic analysis of *B. abortus* TCS protein interactions, Hallez and colleagues previously reported the surprising observation that EssR (then, generically annotated as TcbR) and CenR interact in a yeast two-hybrid assay (68). EssR and CenR were the only two *B. abortus* RRs that showed interaction in their genomescale screen. To test EssR and CenR protein-protein interaction in a heterologous system, we used a bacterial two-hybrid approach based on the T18 and T25 domains of the split adenylate cyclase enzyme (69). CenR and EssR showed strong interactions when fused to either adenylate cyclase domain, while the homomeric CenR-CenR and EssR-EssR combinations showed no evidence of interaction (Figure 5A).

**Figure 5:**
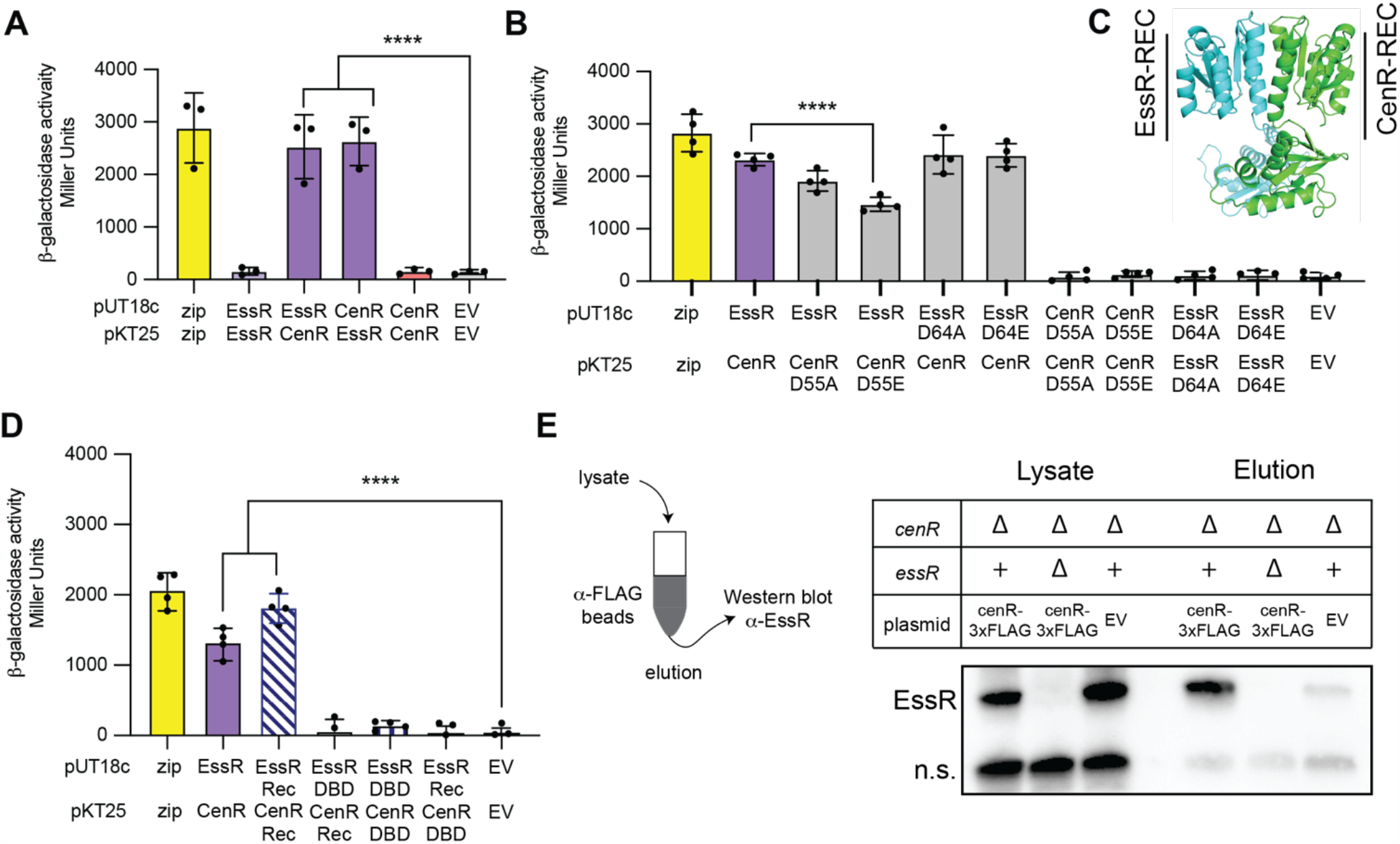
CenR and EssR physically interact in a heterologous system and in *B. ovis*. (A,B) Measurement of homomeric and heteromeric interactions between EssR and CenR (and their aspartyl phosphorylation site mutants; D®A & D®E) using an *E. coli* bacterial two-hybrid assay. Proteins were fused to split adenylate cyclase fragments in vectors pUT18c and pKT25. Positive control (zip) and empty vector (EV) negative controls are shown. (C) CenR:EssR heterodimer structure showing interaction at the α4-β5-α5 structural face of each protein as predicted by AFComplex2 (71); CenR (green) and EssR (blue) (D) Test for interactions between DNA-binding domain (DBD) and receiver domain (Rec) fragments of EssR and CenR by bacterial two-hybrid. Error bars represent the standard deviation of 4 biological replicates. Statistical significance was calculated using one-way ANOVA, followed by Dunnett’s multiple comparisons test (p < 0.0005, ***; p < 0.0001, ****) to EV control or WT CenR-EssR interaction. (E) Co-immunoprecipitation of EssR and CenR-3xFLAG from *B. ovis* lysate. (left) CenR-3xFLAG was captured using anti-FLAG beads, which were washed before elution. (right) EssR association with CenR-3xFLAG in the eluate was monitored by Western blot using polyclonal antiserum to EssR (a-EssR). Non-specific (n.s.) cross-reactive band is shown as an indicator of loading. Representative blot from three biological replicates.

To test the contribution of the conserved aspartyl phosphorylation sites of CenR and EssR to the observed two-hybrid interaction, we fused putative phosphomimetic (DàE) or non-phosphorylatable (DàA) alleles of either CenR or EssR to the T18 and T25 fragments of adenylate cyclase. Both EssR_D64A_ and EssR_D64E_ interacted with CenR to the same extent as WT EssR (Figure 5B). CenR_D55E_ had significantly reduced interaction with EssR, while CenR_D55A_ interaction with EssR was not significantly different. These results provide evidence for a model in which CenR phosphorylation attenuates its interaction with EssR. Homomeric CenR-CenR or EssR-EssR interactions were again not observed in our two-hybrid assay for either the DàA or DàE mutants (Figure 5B).

We used the neural network models of AlphaFold2 (70), as implemented in AF2Complex (71), to develop hypotheses about the structural basis of CenR-EssR interaction. The results of this computation predicted that these two response regulators interact primarily through their receiver (REC) domains rather than their DNA-binding (DBD) domains (Figure 5C). Predicted (1:1) CenR:EssR complex structures showed parallel REC domain heterodimers with significant buried surface area in a region of similar primary structure of CenR and EssR corresponding to the α4-β5-α5 structural face of each protein (Figure S1); this is a well-established REC domain interaction interface (72). To test if CenR and EssR interact via their REC domains, we again used a bacterial two-hybrid assay. The measured interaction of the isolated CenR and EssR REC domains was comparable to that of the full-length proteins (Figure 5D). None of the other DBD-DBD or DBD-REC combinations had significant interactions.

### CenR and EssR interact in *B. ovis*

Given that CenR and EssR strongly interact via their REC domains in a heterologous system, we sought to test whether these two proteins interact in *B. ovis* cells by co-immunoprecipitation. Briefly, we expressed a *cenR-3xFLAG* fusion from the native *cenR* promoter on a lowcopy plasmid in strains lacking either endogenous CenR (*ΔcenR*) or lacking both response regulators (*ΔcenR ΔessR*) and applied the crosslinked lysates to anti-FLAG magnetic beads. After multiple washing steps we eluted bound protein, reversed the crosslinks, and resolved the eluate by SDS-PAGE. Western blot using polyclonal EssR antiserum revealed clear EssR bands in the lysates of both *ΔcenR / cenR-3xFLAG* and *ΔcenR* vector control strains, but not in *ΔcenR ΔessR / cenR-3xFLAG*. Among the three eluate fractions, only the strain expressing both *cenR-3xFLAG* and endogenous *essR* yielded a strong EssR band on the Western blot (Figure 5E). These results provide evidence that the CenR-EssR interaction demonstrated by bacterial two-hybrid assay occurs in *B. ovis* cells.

### CenR enhances the rate of phosphoryl transfer from EssS to EssR

The related phenotypes of *cenR* and *essR* mutants and the fact that these two response regulators physically interact in cells raised questions about the biochemical consequence(s) of EssR/CenR interaction on histidine kinase autophosphorylation and phosphoryl transfer. The data presented to this point do not clearly establish the identity of a sensor kinase in this system, though we considered EssS to be a strong candidate based on genomic proximity of *essS* and *essR*, and largely overlapping phenotypes between Δ*essS* and Δ*essR* strains. To test this hypothesis, the cytoplasmic kinase domain of EssS (aa 191-488) and full-length EssR and CenR response regulators were purified. The EssS kinase domain autophosphorylated *in vitro*; incubating EssS kinase domain with EssR resulted in rapid loss of phospho-EssS (EssS∼P) signal and a concomitant increase in phospho-EssR (EssR∼P) signal within 30s, indicating phosphoryl transfer (Figure 6A). Incubation of EssS with CenR resulted in no detectable phosphoryl transfer by 600s, even when CenR was added in 50 molar excess. We conclude that EssS is the cognate kinase for EssR, and that EssS does not directly phosphorylate CenR.

**Figure 6:**
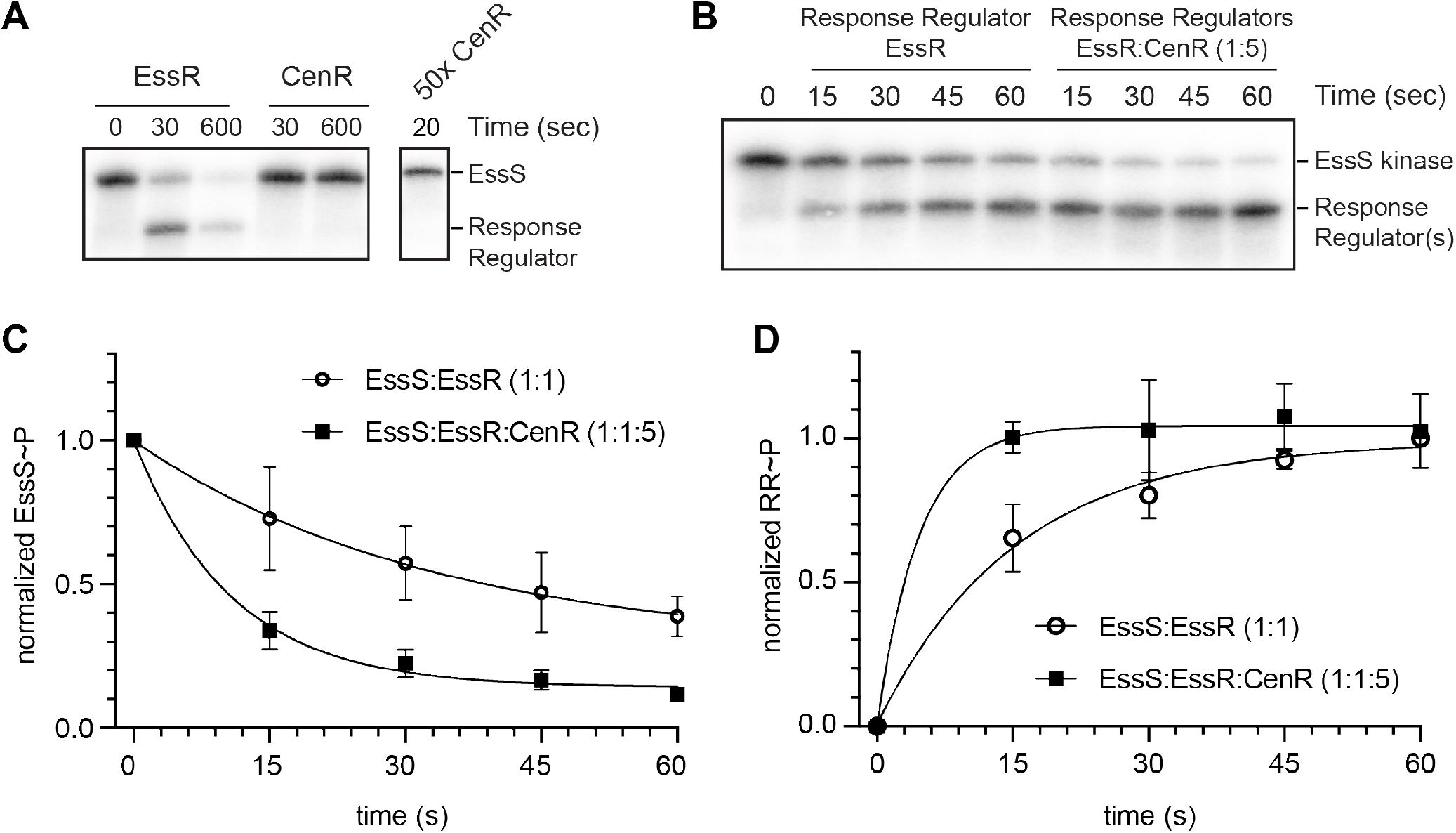
EssS specifically phosphorylates EssR; CenR stimulates phosphoryl transfer from EssS to EssR. (A) EssS autophosphorylation and phosphoryl transfer assay. Phosphoryl transfer from EssS to either EssR or CenR was assessed at 30s and 600s timepoints. 50x molar excess of CenR was added to phospho-EssS to test kinase specificity. Lower molecular weight band corresponds to phospho-EssR. B) Phosphoryl transfer kinetics from EssS to EssR at a 1:1 molar ratio and at a 1:1:5 (EssS:EssR:CenR) molar ratio. C) Normalized EssS-P dephosphorylation kinetics at a 1:1 molar ratio of EssS to EssR and at a 1:1:5 (EssS:EssR:CenR) ratio. D) Normalized EssR phosphorylation kinetics at the same molar ratios as panel C. Points and error bars represent the mean ± standard deviation of three replicates.

Though EssS does not phosphorylate CenR *in vitro*, we postulated that CenR may influence activity of the EssS- EssR TCS. To test this idea, we first mixed equimolar EssS and EssR with increasing concentrations of CenR. Supplementing an EssS-EssR reaction mixture with CenR at a 1:1:1 molar ratio resulted in a 20% increase in EssS∼P dephosphorylation after 20s relative to a 1:1 EssS-EssR reaction (Figure S5). Adding CenR to the reaction mixture at 5x molar excess (1:1:5), further enhanced EssS∼P dephosphorylation (Figure S5). The reduction in EssS∼P upon addition of CenR coincides with an increase in RR∼P. The effect of CenR addition on EssS∼P levels (at 20 s) saturated at a 1:1:5 ratio (Figure S5). Taken together, the data indicate that phosphoryl transfer from EssS to EssR is accelerated by addition of CenR. To further investigate the effect of CenR on the kinetics of phosphoryl transfer between EssS and EssR, we measured both EssS∼P and EssR∼P signal over a 1-minute time course. EssS to EssR phosphoryl transfer reactions containing 5-molar excess CenR had an enhanced rate of (apparent) EssR∼P production and an enhanced rate of EssS∼P loss compared to reactions with EssS and EssR only (Figure 6B-D); these experiments do not rule out the possibility that CenR is phosphorylated by EssS when EssR is present as EssR and CenR have similar molecular weights. In the absence of CenR, we observed maximal EssR∼P signal after 45-60s. When CenR was present, levels of the band we attribute to EssR∼P were maximal by 15s. These results provide evidence that CenR stimulates phosphoryl transfer from EssS to EssR.

### EssR and CenR determine protein levels of each other via a post-transcriptional mechanism

Stimulation of phosphoryl transfer from EssS to EssR by the non-cognate CenR protein provides one explanation for the related phenotypes of the Δ*cenR, ΔessS* and *ΔessR* mutants. We sought to test whether the CenR protein may have other regulatory effects on the cognate EssS-EssR TCS pair in *B. ovis*. EssR and CenR do not regulate each other’s transcription (Table S3) but do physically interact (Figure 5). We hypothesized that CenR interaction with EssR could impact steady-state EssR protein levels through a post-transcriptional mechanism, and vice versa. To test this hypothesis, we measured EssR protein by Western blot in WT and Δ*cenR*. Deletion of *cenR* resulted in a significant reduction (∼70%) of EssR protein levels. The Δ*essS* strain also had significantly reduced EssR levels, though the effect was not as large as Δ*cenR* (Figure 7). The impact of EssS on EssR protein levels is not apparently a consequence of phosphorylation as steady-state levels of EssR, EssR(D64A) and EssR(D64E) did not differ significantly. The mechanism by which CenR affects EssR protein levels is not known, but reduced EssR levels in a Δ*cenR* background provide an additional explanation of the phenotypic congruence of the Δ*cenR* and Δ*essR* strains. The intermediate level of EssR protein in the *ΔessS* strain is consistent with the intermediate phenotype of this mutant in plate stress assays. We further tested whether CenR protein levels are impacted by *essR* by measuring CenR-3xFLAG in *ΔcenR / cenR-3xFLAG* and *ΔcenR ΔessR / cenR-3xFLAG* strains. CenR-3xFLAG was »60% lower in a strain lacking *essR*. We conclude that CenR and EssR regulate each other’s protein levels through a post-transcriptional mechanism.

**Figure 7:**
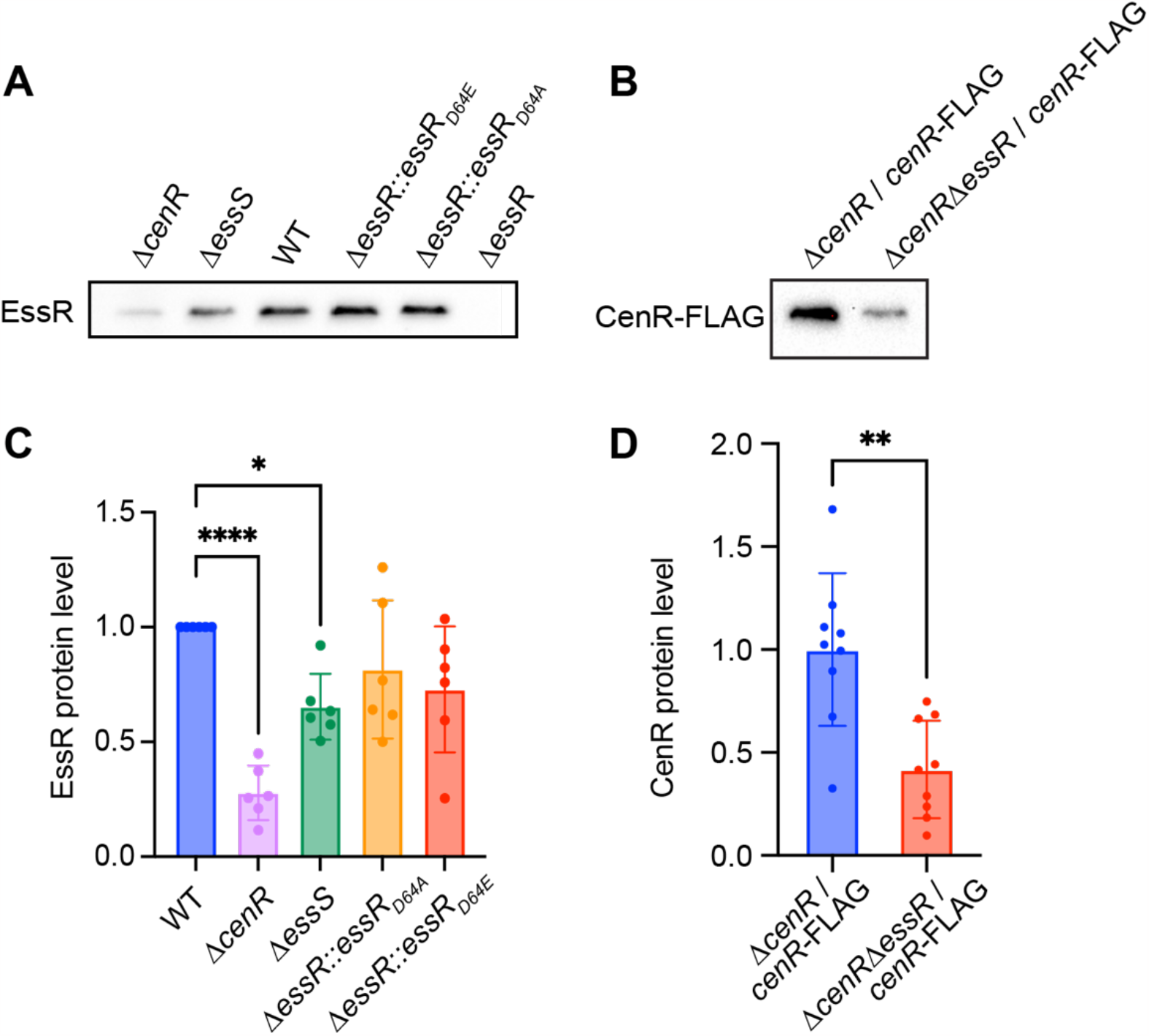
CenR and EssR proteins levels are influenced by each other. (A) a-EssR Western blot of lysate from WT *B. ovis* and *B. ovis* expressing EssR(D64A), EssR(D64E), and in strains lacking *essS* (D*essS*), *cenR* (D*cenR*) or *essR (ΔessR*). (B) a-FLAG Western blot of lysate from *B. ovis* strains lacking *cenR* (*ΔcenR*) or *cenR* and *essR (ΔcenR ΔessR*) and expressing *cenR-3xFLAG* from pQF plasmid. (C) Quantification of EssR band intensities normalized to the EssR intensity from WT on each blot. Error bars represent the mean ± standard deviation of six replicates. Statistical significance was calculated using one-way ANOVA, followed by Dunnett’s multiple comparisons test to WT (p < 0.05, *; p < 0.0001, ****). (D) Quantification of CenR-3xFLAG band intensities normalized to the average intensity from *ΔcenR / cenR-3xFLAG* on each blot. Error bars represent the mean ± standard deviation of nine samples assayed on three independent blots. Statistical significance was calculated using an unpaired t test (p < 0.005, **).

## Discussion

Multiple TCS genes are typically present in bacterial genomes, and the proteins they encode most often function separately to regulate distinct transcriptional responses (20). Our systematic analysis of *B. ovis* TCS genes revealed two DNA-binding response regulators, *cenR* and *essR*, that had related morphological, stress resistance, and infection phenotypes when deleted. These results informed the hypothesis that CenR and EssR work together to execute their functions in the cell and led to the discovery of a new cell envelope regulatory system (EssS-EssR) and a new mechanism of TCS regulation in bacteria.

### Discovery of a cell envelope regulatory system in *Brucella*

TCS proteins play a key role in the regulation of cell envelope biogenesis and homeostasis in the bacterial kingdom (29, 73, 74). The function of the EssR-EssS TCS protein pair had not been defined in any species prior to this study, though this system is conserved in many *Alphaproteobacteria*. Our data provide evidence that these two proteins play an important role in *Brucella* resistance to cell envelope disruptors *in vitro*, and in regulating processes important for intracellular replication in a macrophage infection model. Apparent orthologs of the sensor kinase, EssS, are present in select genera across the orders *Hyphomicrobiales, Caulobacterales, Rhodobacterales, Rhodospirillales*, and *Rickettsiales*; EssR has a similar phylogenetic distribution (Figure S6). The functional importance of EssR-EssS in *Alphaproteobacteria* is evidenced by the fact that it is 1 of only 5 TCS signaling pairs (NtrXY, PhoBR, RegBA, ChvGI/BvgRS, & EssRS) in the highly streamlined SAR11 genome (*Pelagibacter ubique*). The HK domain of EssS has low sequence identity to other well-studied Gramnegative envelope regulators (e.g. CpxA and EnvZ), but multiple sequence alignment models in the conserved domain database (CDD) (52) suggest that the EssS, CpxA, and EnvZ HKs have common ancestry (e-value < 10^−40^). Likewise, EssR is most closely related to the OmpR sequence family in the CDD (e-value < 10^−70^). OmpR functions as the cognate regulator of the EnvZ kinase in enteric bacteria (53).

EssS and EssR clearly form a cognate signaling pair *in vitro* as evidenced by specific phosphoryl transfer from the EssS kinase domain to EssR on a fast time scale (Figure 6). However, the phenotypes of the *ΔessS* and *ΔessR* strains are not equivalent in an *in vitro* model of cell envelope stress. Under most challenges (SDS, carbenicillin, EDTA, and polymyxin B) the defect of *ΔessR* was more severe than *ΔessS* (Figure 1, S1 and S2), and these mutants have opposite phenotypes when exposed to high NaCl: the *essR* mutant is NaCl resistant while the *essS* mutant is sensitive compared to WT (Figure S2). The mechanism underlying the opposing NaCl phenotypes of these strains merits further investigation. The phenotypes of *ΔessS* and *ΔessR* are equivalent in a macrophage infection model (Figure 3). This result provides evidence that EssS-dependent phosphorylation (or dephosphorylation) of EssR is more important for system function in the complex environment of the intracellular niche than it is in a simple *in vitro* agar plate assay.

### An unexpected functional role for the conserved cell envelope regulator, CenR

The result that *B. ovis* CenR confers resistance to SDS and carbenicillin (Figures 1 & 2) was not unexpected considering the phenotypes of *cenR* mutants in other Alphaproteobacteria. CenR was first described as an essential RR in *C. crescentus*, where it functions to regulate cell envelope structure (34), and is now known to be conserved in many alphaproteobacterial orders (33). Recent work has shown that *cenR* is essential in *R. sphaeroides*, where it controls transcription of the Tol-Pal outer membrane complex and other cell envelope genes (33), and in *Sinorhizobium meliloti* where it mediates osmotolerance and oxidative stress resistance (32). In all three of these species, CenR is regulated by a cognate histidine kinase, CenK. Our data show that *cenR* is not essential for *B. ovis* growth or division under standard culture conditions and indicate that it may be an orphan response regulator, which is consistent with a previous report in *B. melitensis* (49). *B. ovis cenR* (and *essR*-*essS*) do not function to mitigate acid stress *in vitro*. Though a *B. melitensis cenR* mutant was previously reported to be acid sensitive (48), the treatment protocol and measured pH range differ substantially between our *B. ovis* study and the *B. melitensis* study. The possibility that a *Brucella* HK phosphorylates (or dephosphorylates) CenR under certain conditions cannot be conclusively ruled out from our data. Indeed, there is some evidence that the conserved CenR aspartyl phosphorylation site can impact CenR-EssR interaction (Figure 5) and replication in the intracellular niche (Figure 3). Nonetheless, both *cenR*_*D55A*_ and *cenR*_*D55E*_ alleles fully complement the agar plate stress phenotypes of D*cenR*. And unlike *R. sphaeroides* and *C. crescentus*, where *cenR* depletion results in major defects in cell envelope structure, the impact of *cenR* deletion on *B. ovis* cell morphology is small: *B. ovis* D*cenR* mutants are slightly (but significantly) larger than WT (Figure S3), but the morphology of the mutant cells otherwise appears normal. These results indicate that CenR function in *Brucella ovis* differs somewhat from *Caulobacter, Rhodobacter*, and *Sinorhizobium*.

### CenR is a post-transcriptional regulator of the EssS- EssR two-component system

Cross-regulation between TCSs is uncommon, though there is experimental support for direct interactions between otherwise distinct TCS HK-RR protein pairs for limited number of systems (20). For example, the NarXL and NarQP systems of *Escherichia coli* cross-phosphorylate to tune nitrate and nitrite respiratory processes (75), and in *C. crescentus*, a consortium of sensor kinases that coordinately regulate cell adhesion in response to a range of environmental cues physically interact in cells (23). In this manuscript, we present evidence for a mode of TCS cross-regulation in which a non-cognate RR (CenR) directly stimulates the phosphoryl transfer activity of a cognate HK-RR protein pair (EssS-EssR) (Figure 6 and Figure 8). We further demonstrate that CenR and EssR reciprocally regulate their protein levels in *B. ovis* cells via a post-transcriptional mechanism (Figure 7). EssR and CenR physically interact via their receiver domains (Figure 5), and it seems most likely that CenR and EssR protect each other from proteolytic degradation in the *Brucella* cell though we cannot rule out other post-transcriptional models at this time. The positive effect of CenR on EssS-EssR phosphoryl transfer activity and the positive effect of EssR and CenR protein levels on each other are consistent with the congruent phenotypes of strains lacking *cenR* and *essR*.

**Figure 8:**
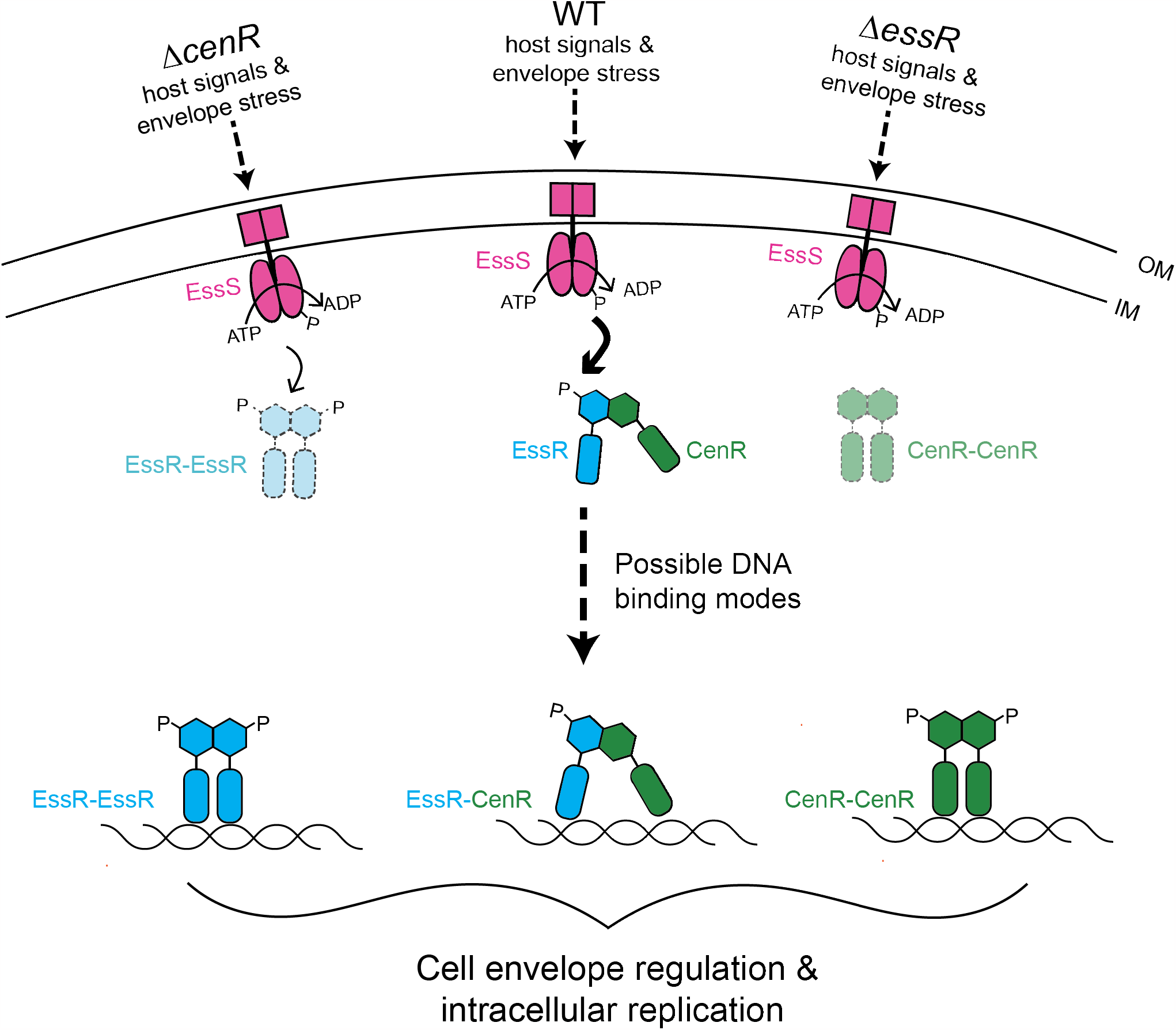
Model of EssS-EssR-CenR-dependent gene regulation in *B. ovis*. Upon detection of host signals, EssS (pink) autophosphorylates and transfers a phosphoryl group (P) to EssR (blue). CenR (green) supports EssS-EssR signal transduction by directly stimulating phosphoryl transfer from EssS and EssR via a receiver domain interaction. Loss of CenR results in diminished EssR levels (light blue with dashed outlines) and loss of EssR results in diminished CenR levels (light green with dashed outlines), via a post-transcriptional mechanism. CenR and EssR directly interact with an overlapping set of promoters, and three possible modes of transcriptional regulation by EssR and CenR are shown: EssR-EssR homodimer, EssR-CenR heterodimer, and/or CenR-CenR homodimers binding to target promoters. Genes regulated by the EssS-EssR-CenR system impact *B. ovis* cell size, contribute to growth/survival during cell envelope stress *in vitro*, and support intracellular replication in a macrophage infection model.

However, CenR does not simply control the activity and levels EssS-EssR. CenR is itself a DNA binding protein, and we have presented evidence that EssR and CenR both directly and indirectly regulate transcription of a highly correlated set of genes that includes multiple transporters and cell wall metabolism genes (Figure 4). These two transcriptional regulators directly bind shared and unique sets of sites on *B. ovis* chromosomes 1 and 2. It is possible that CenR and EssR bind DNA as heterodimers, which has been described for the BldM and WhiI response regulators of *Streptomyces* (76), and the RcsB regulator of *E. coli* with GadE (77) and BglJ (78). These heterodimeric regulators are competent to control different classes of promoters depending on oligomeric state; a similar mechanism may exist for EssR and CenR though we have not identified distinct promoter classes in our data. It is plausible that CenR and EssR bind as homodimers and heterodimers considering the pattern of unique and overlapping genes/promoters in the transcriptomic and ChIP-seq datasets. Future studies aimed at deciphering molecular features of environmental signal detection by the EssS sensor kinase, allosteric regulation of TCS activity by CenR, and transcriptional control by the CenR and EssR regulators will generally inform our understanding of the evolution of cell envelope regulatory systems in bacteria. More specifically, investigation of these proteins will illuminate mechanisms by which *Brucella* replicate in the intracellular niche and spread from cell to cell in face of harsh immune stresses encountered within the host.

## Materials and Methods

### Bacterial strains and growth conditions

All *Brucella ovis* ATCC25840 and derivative strains were grown on Tryptic soy agar (BD Difco) + 5% sheep blood (Quad Five) (TSAB) or in Brucella broth at 37°C with 5% CO_2_ supplementation. Growth medium was supplemented with 50 µg/ml kanamycin, 30 µg/ml oxytetracycline, 2 µg/ml carbenicillin, 0.004%-0.0045% sodium dodecyl sulfate (SDS), 2.75 mM EDTA, or 215 mM NaCl when necessary.

All *Escherichia coli* strains were grown in lysogeny broth (LB) or on LB solidified with 1.5% w/v agar. *E. coli* Top10 and WM3064 strains were incubated at 37°C and BTH101 strains were incubated at 30°C. WM3064 was grown with 30 µM diaminopimelic acid (DAP) supplementation. Medium was supplemented with 50 µg/ml kanamycin, 12 µg/ml oxytetracycline or 100 µg/ml carbenicillin when necessary. Primer, plasmid, and strain information are available in Table S5.

### Essential gene calculations using *B*. *ovis* Tn-himar sequencing data

We constructed a library of *B. ovis* transposon mutants as described previously (15). Briefly, the *E. coli* strain APA752, harboring the pKMW3 mariner transposon library, was conjugated into WT *B. ovis* and transposon bearing strains were selected on TSAB supplemented with 50 µg/ml kanamycin. Following an outgrowth of pooled mutants in 250 ml Brucella broth to OD_600_ ≈ 0.6, cells were frozen in 25% glycerol in 1 ml aliquots. An aliquot was thawed for gDNA extraction and subsequent insertion site mapping, as described (79). The library contained insertions at over 50,000 unique sites in the genome. Insertion site sequencing data is available in the NCBI sequence read archive under accession SRR19632676. Using the insertion site mapping data from this Tn-seq dataset, we applied the HMM and Gumbel algorithms in the TRANSIT (37) software package to identify candidate essential genes (see Table S1).

### Chromosomal deletion strain construction

The double-crossover recombination method was used to generate all *B. ovis* strains bearing in-frame, unmarked gene deletions. Approximately 500 base pair (bp) upstream or downstream of the target gene, including 9- 120 bp of the 3’ and 5’ ends of the target gene, were amplified by PCR using KOD Xtreme polymerase (Novagen) using primers (Table S5) specific to these regions and the *B. ovis* genomic DNA as template. The DNA fragments were then inserted into the *sacB-* containing suicide plasmid pNPTS138 either through Gibson assembly or restriction enzyme digestion and ligation. The ligated plasmids were first chemically transformed into competent *E. coli* Top10. After sequencing confirmation, plasmids were transformed into chemically competent *E. coli* WM3064 (strain originally produced by W. Metcalf), a DAP auxotroph conjugation-competent donor strain. Plasmids were transferred to *B. ovis* through conjugation. Primary recombinants were selected on TSAB supplemented with kanamycin. After outgrowth in non-selective growth in Brucella broth for 8 hours, clones in which a second recombination removed the plasmid were identified through counter selection with 5% sucrose. Colony PCR was performed on kanamycin-sensitive colonies to distinguish clones bearing the deletion allele from those bearing the WT allele.

### Complementation strain construction

To engineer genetic complementation constructs, target genes were amplified by KOD polymerase, including ∼300 bp upstream and ∼50 bp downstream of each target gene. The PCR products were purified and inserted into plasmid pUC18-mTn7 by restriction enzyme digestion and ligation, followed by chemical transformation into *E. coli* TOP10 cells. After sequence confirmation, the mTn7 plasmids were transformed into chemically competent *E. coli* WM3064. These plasmids were co-conjugated into *B. ovis* strains with a Tn7 integrase expressing suicide helper plasmid, pTNS3, which is also carried by WM3064. *B. ovis* colonies carrying the integrated mTn7 constructs at the *glmS* locus were selected on TSAB containing kanamycin.

### Agar plate growth/stress assays

After 2 days of growth on TSAB or TSAB supplemented with kanamycin, *B. ovis* cells were collected and resuspended in sterile phosphate-buffered saline (PBS) to an OD_600_ = 0.3. Each strain was serially diluted in PBS using a 10^−1^ dilution factor. 5 µl of each dilution was plated onto either TSAB, TSAB containing 0.004%-0.0045% SDS, 2 µg/ml carbenicillin, 2.75 mM EDTA, or 215 mM NaCl. After 3 days of incubation for TSAB, 4 days for TSAB + 0.004%-0.0045% SDS and 5 days for TSAB + 2 µg/ml carbenicillin, 2.75 mM EDTA, and 215 mM NaCl, growth was documented photographically.

### Polymyxin B stress assay

After 2 days of growth on TSAB supplemented with kanamycin, *B. ovis* cells were collected and resuspended in 1 mL of Brucella broth at an OD_600_ =0.3. Cells were split and one portion was treated with 1 mg/mL polymyxin B and the other was untreated. Both treated and untreated groups were incubated at 37°C with 5% CO_2_ supplementation for 80 minutes. Each culture was then 10-fold serially diluted in PBS and 5 µl of each dilution was spotted onto TSAB. After 3 days of incubation for untreated group and 4 days for treated group, growth was documented photographically.

### Macrophage infection assay

THP-1 cells were grown in Roswell Park Memorial Institute Medium (RPMI) + 10% heat inactivated fetal bovine serum (FBS) at 37°C with 5% CO_2_ supplementation. Three days prior to infection, THP-1 cells were seeded in a 96-well plate at a concentration of 10^6^ cells/ml in 100 µl/well of fresh RPMI + 10% FBS with an addition of 50ng/ml phorbol 12-myristate-13-acetate (PMA) to induce differentiation. After three days at 37°C in 5% CO_2,_ *B. ovis* cells were resuspended at a concentration of 10^8^ CFU/ml (OD_600_ = 0.15) in RPMI + 10% FBS and added to THP-1 cells at a multiplicity of infection (MOI) of 100. The 96-well plate containing *B. ovis* and THP-1 cells was centrifuged at 150 x g at room temperature for 5 minutes and incubated at 37°C with 5% CO_2_ for 1h. Media was removed and fresh media containing 50 µg/ml gentamicin was added to kill extracellular *B. ovis* that were not internalized. The plate was incubated for another hour at 37°C with 5% CO_2_.

Media was removed and fresh media containing 25 µg/ml gentamicin was added to each well except for 2h-p.i. wells. A 2h, 6h, 24h, and 48h post-infection, *B. ovis* were enumerated by removing the media and washing the cells with PBS once and incubating at 37°C for 10 minutes. Then PBS was removed and 200 µl dH_2_O was added to each well and the plate was incubated at room temperature for 10 minutes to lyse the THP-1 cells. *B. ovis* cells were collected and serial diluted with a 10^−1^ dilution factor in PBS and plated on TSAB. Plates were incubated for 3 days and CFU were enumerated.

### Acid stress assay

As previously described (7), *Brucella* cells were grown on TSAB for 2 days before being inoculated into 5 mL Brucella broth and shaken for 24h. OD_600_ was adjusted to 1.5 and 50 µl of cells were added to either 5 mL plain Brucella broth or 5 mL Brucella broth pH 4.2 (final OD_600_ = 0.015). After 2h of shaking incubation, cells were serially diluted and plated onto plain TSAB. After 3 days of incubation, CFU/ml were enumerated.

### Phase contrast microscopy

Samples of *B. ovis* cells were grown on TSAB supplemented with kanamycin and incubated for 2 days. Cells were resuspended in milliQ H_2_O and 1 µl of cells were spotted on an agarose pad on a cover slide and imaged on a Leica DMI 6000 microscope in phase contrast with an HC PL APO 63x/1.4 numeric aperture (NA) oil Ph3 CS2 objective. Images were captured with an Orca-ER digital camera (Hamamatsu) controlled by Leica Application Suite X (Leica). Cell areas were measured from phase contrast images using BacStalk (80).

### Cryo-electron microscopy

*B. ovis* cells were grown on TSAB for 2 days and resuspended in PBS to a OD_600_ ≈ 1. Five µl of cells were placed on a glow-discharged R 3.51 grid (Quantfoil), blotted for 3.5 s, air dried for 20 s, and plunged into liquid ethane using the Vitrobot robotic plunge freezer (Thermo). Samples were stored in liquid nitrogen before imaging. Cells were imaged on a ThermoScientific Talos Arctica Cryo-EM (Thermo) with a -10.00 µm defocus and 17,500x magnification. Images were captured with a Ceta camera (Thermo) with 2.0 s exposure time.

### RNA extraction and sequencing

*B. ovis* strains were grown in Brucella broth at 37°C in 5% CO_2_ overnight. The next day, cultures were back diluted to OD_600_ = 0.05 in 12 ml Brucella broth and incubated on a rotor at 37°C with 5% CO_2_ for 7h. 9 ml of each culture was harvested by centrifugation, and the pellets were immediately resuspended in 1 ml TRIzol. Samples were stored at -80°C until RNA extraction. To extract the RNA, samples were thawed and incubated at 65°C for 10 minutes. 200 µl of chloroform was added to each sample and mixed by vortexing for 15 seconds. Samples were then incubated at room temperature for 5 minutes. Phases were separated by centrifugation at 17,000 x g for 15 minutes at 4°C. The aqueous layer was transferred into a fresh tube. Sample was mixed with 500 ml of isopropanol and stored at -20°C for 2h. After thawing, samples were centrifuged at 13,000 x g for 30 minutes at 4°C to pellet the RNA. Supernatants were discarded and the pellets were washed with 1 ml of ice cold 70% ethanol. Samples were centrifuged at 13,000 x g at 4°C for 5 minutes. Supernatants were discarded and pellets were air dried. Pellets were resuspended in 100 µl RNAse-free H_2_O and incubated at 60°C for 10 minutes. RNA samples were DNase treated using RNeasy Mini kit (Qiagen). RNA samples were sequenced at Microbial Genome Sequencing Center (Pittsburgh, PA) on an Illumina NextSeq 2000 (Illumina). RNA libraries for RNA-seq were prepared using Illumina Stranded RNA library preparation with RiboZero Plus rRNA depletion.

### RNA-seq analysis

All RNA-seq analysis was conducted in CLC Genomics Workbench 21.0.4 (Qiagen). Reads were mapped to reference genome (*Brucella ovis* ATCC 25840, Genbank accessions NC_009504 and NC_009505). Differential expression for RNA-seq default settings were used to calculate the fold change of all annotated genes in mutant strains versus WT. Raw and processed RNA-seq data are publicly available in the NCBI GEO database at accession GSE229183.

### ChIP DNA extraction and sequencing

To build the constructs for ChIP-seq analysis of strains expressing CenR_D55E_ and EssR_D64E_, regions containing each gene (including ∼300 bp upstream) were PCR amplified from the *B. ovis cenR*_D55E_ and *essR*_D64E_ mutant strains with KOD polymerase. Amplified fragments were purified and inserted into plasmid pQF through restriction enzyme digestion and ligation to yield a 3xFLAG fusion at the 3’ end of each gene. This C-terminal FLAG fusion plasmid construct was conjugated into *B. ovis* mutants using WM3064 as described above.

Cells were grown on TSAB plates supplemented with 30 mg/ml oxytetracycline for three days and resuspended in Brucella broth. Formaldehyde was added to a final concentration of 1% (w/v) to crosslink, and samples were incubated for 10 minutes on shaker. Crosslinker was quenched by adding 0.125 M glycine and was shaken for an additional 5 minutes. Cells were then centrifuged at 7,196 x g for 5 minutes at 4°C and pellets were washed 4 times with cold PBS pH 7.5 and resuspended in 900 µl buffer [10 mM Tris pH 8, 1 mM EDTA, protease inhibitor cocktail tablet (Roche)]. 100 µl of 10 mg/ml lysozyme was added to each sample, which were then vortexed and incubated at 37°C for 30 minutes before adding a final concentration of 0.1% SDS (w/v) to each sample. The samples were then sonicated on ice for 15 cycles (20% magnitude, 20 seconds on/off) and centrifuged at 15,000xg for 10 minutes at 4°C. Supernatants were collected, Triton X-100 was added to a concentration of 1%, and lysates were added to 30 µl SureBead Protein A magnetic agarose beads (BioRad) equlibrated with binding buffer [10 mM Tris pH 8, 1 mM EDTA pH8, 0.1% SDS, 1% Triton X-100] and incubated for 30 minutes at room temperature; this step has empirically improved signal for our FLAG IP protocols. Beads were collected on a magnetic stand, the supernatant was transferred to a fresh tube, and 5% of each sample supernatant was removed as the input DNA samples. 100 µl of α-FLAG magnetic agarose beads were then washed three times with binding buffer and incubated overnight at 4°C shaking in 500 µl binding buffer plus 1% BSA to equilibrate the beads. The following day, α-FLAG beads were washed four times in binding buffer and cell lysates were added to the pre-washed beads and gently vortexed. Samples were incubated for 3h at room temperature with mixing and α-FLAG beads were collected on a magnetic stand and serially washed in 500 µl low-salt buffer [50 mM HEPES pH7.5, 1% Triton X-100, 150 mM NaCl], 500 µl high-salt buffer [50mM HEPES pH 7.5, 1% Triton X-100, 500 mM NaCl], and 500 µl LiCl buffer [10 mM Tris pH 8, 1 mM EDTA pH 8, 1% Triton X- 100, 0.5% IGEPAL^®^ CA-630, 150 mM LiCl]. To elute the protein-DNA complex bound to the beads, 100 µl elution buffer [10 mM Tris pH 8.0, 1 mM EDTA pH 8.0, 1% SDS, 100 ng/µl 3x FLAG peptide] was added to the beads and incubated at room temperature for 30 minutes with mixing. The eluate was collected, and the elution step was repeated. Input samples were brought to the same volume as output samples with elution buffer. For RNase A treatment, a final concentration of 300 mM NaCl and 100 µg/ml RNase A were added to each input and output sample and incubated at 37°C for 30 minutes. Proteinase K was added to a final concentration of 200 µg/ml and the samples were incubated overnight at 65°C to reverse crosslinks. ChIP DNA was purified with Zymo ChIP DNA Clean & Concentrator kit. The ChIP-seq library was prepared using Illumina DNA prep kit and IDT 10bp UDI indices; DNA samples were sequenced at SeqCenter (Pittsburgh, PA) on an Illumina Nextseq 2000.

### ChIP-seq analysis

Output and input DNA sequences were first mapped to the reference genome (*Brucella ovis* ATCC 25840, Genbank accessions NC_009504 and NC_009505) in Galaxy using bowtie2 (81). ChIP-seq enriched peak calls from the mapping output data were carried out in Genrich (https://github.com/jsh58/Genrich) with parameters maximum q-value = 0.05, and minimum AUC (area under the curve) = 20; PCR duplicates were removed. bamCoverage (82) was used to create bigwig file tracks for each replicate, and peaks were visualized in IGB (83). Raw and processed ChIP-seq data are publicly available in the NCBI GEO database at accession GSE229183.

### Bacterial two hybrid β-galactosidase assay

The DNA fragments of the full-length, point mutant, and specific domains of *cenR* and *essR* were PCR amplified by KOD polymerase and cloned into split adenylyl cyclase plasmids (either pKT25 or pUT18c) (69) through restriction digestion and ligation. Each pair of pKT25 and pUT18c plasmids were co-transformed into chemically competent *E. coli* BTH101 cells. BTH101 strains carrying both pKT25 and pUT18c plasmids were grown in LB and incubated at 30°C overnight while shaking. Fresh LB + 500 µM IPTG was inoculated with 100 µl of overnight culture and incubated at 30°C until OD_600_ ≈ 0.3-0.4. To assess two-hybrid interaction via reconstitution of adenylyl cyclase activity, 100 µl of culture was mixed with 100 µl chloroform and vortexed vigorously. 700 µl Z buffer (0.06 M Na_2_HPO_4_, 0.04 M NaH_2_PO_4_, 0.01 M KCl, 0.001 M MgSO_4_) and 200 µl ortho-nitrophenyl- β-galactoside (ONPG) was added to each sample. The reactions were stopped with 1 ml of Na_2_CO_3_ after 7.8 minutes and colorimetric conversion of ONPG was measured at 420 nm (A_420_) was measured. Miller Units were calculated as MU = A_420_ × 1000 / (OD_600_ x time x volume of cells).

### Structure prediction

The heterodimeric structure of CenR and EssR was predicted using the protein complex prediction package AF2Complex (71), which utilizes the neural network models of AlphaFold2 (70). Computations were carried out on the Michigan State University high-performance computing cluster.

### EssR-CenR co-immunoprecipitation assay

Strain construction, lysate production, and α-FLAG magnetic agarose beads preparation were the same as described above for ChIP-seq sample preparation. After preparation and centrifugation of the lysates, the supernatants were collected and 50 µl of each sample was mixed with equal amount of SDS loading buffer and stored at -20°C. For FLAG IP, remaining lysates were applied to pre-washed α-FLAG beads and incubated at room temperature for 3h with shaking. Beads were washed and eluted as described above, eluate from each sample was incubated at 65°C overnight to reverse crosslinking, and samples were then mixed with equal volume of SDS loading buffer. All samples collected were heated to 95°C for 5 minutes before resolving on a 12% mini-PROTEAN precast gel (BioRad). A Western blot using polyclonal EssR antiserum was conducted following the protocol outlined in the Western blot method section below. The membrane was imaged using BioRad Chemi- Doc Imaging System (BioRad).

### Protein purification

DNA fragments that encode full length CenR and EssR, and EssS residues 191-488 (EssS for short) were PCR amplified by KOD polymerase and inserted into a pET23b-His6-SUMO expression vector through restriction digestion and ligation. Vectors were transformed into chemically competent *E. coli* BL21 Rosetta (DE3) / pLysS. All strains were grown in LB medium at 37°C and induced with 0.5 mM isopropyl β-D- 1-thiogalactopyranoside (IPTG) at OD_600_ ≈ 0.5. Cell pellets were harvested after 2h of induction at 37°C and stored at -80°C until purification.

For protein purification, cell pellets were resuspended in 25 ml of lysis buffer (25 mM Tris pH8, 125 mM NaCl, 10 mM imidazole) with the addition of 1mM phenylmethylsulfonyl fluoride (PMSF) and 5 µg/ml DNase I. Samples were lysed through sonication on ice (20% magnitude, 20 seconds on/off) until the lysates were clear. Lysates were centrifuged, and the supernatant was added to 3 ml of Ni-nitrilotriacetic acid (NTA) superflow resin (Qiagen) and the mixture was applied onto a gravity drip column. The resin was washed with 20 ml of lysis buffer, 50 ml of wash buffer (25 mM Tris pH8, 150 mM NaCl, 30 mM imidazole) and the proteins were collected in 20 ml of elution buffer (25 mM Tris pH8, 150 mM NaCl, 300 mM imidazole) at 4°C. For CenR and EssR, the elution was dialyzed and the His6-SUMO tags were cleaved with ubiquitin-like-specific protease 1 (Ulp1) overnight at 4°C in dialysis buffer (25 mM Tris pH8, 150 mM NaCl). The next day, the digested protein was mixed with NTA superflow resin. The mixture was applied onto a gravity column and the purified and cleaved protein was collected from the flow through. His6-SUMO-EssS was dialyzed in dialysis buffer overnight at 4°C and collected. All proteins were stored at 4°C.

### *In vitro* EssS-EssR phosphoryl transfer assay

Purified His-SUMO-EssS(191-488), CenR, and EssR were diluted to 5 µM with 1x kinase buffer (250 mM Tris pH8, 50 mM KCl, 10 mM MgCl_2_, 10 mM CaCl_2_, 1 mM DTT). 5 µl of ATP mix (4.5 µl 250 µM ATP, pH 7, 0.5 µl [32P]-ATP (10 µCi/µl) was added to 45 µl of 5 µM His-SUMO-EssS and the protein was allowed to autophosphorylate for 1h at room temperature. 5 µl of His-SUMO-EssS∼P was mixed with 5 µl 1x kinase buffer and SDS loading buffer as the autophosphorylation control. To assess phosphoryl transfer, His-SUMO-EssS∼P was mixed with equimolar of CenR, EssR, or a CenR/EssR mixture at various CenR:EssR molar ratios. Each phosphoryl transfer reaction was quenched with equal volume of SDS loading buffer. Reaction samples were loaded onto a 12% mini-PROTEAN precast gel (BioRad) and resolved at 180V at room temperature. The dye front of the gel was cut off and the rest of the gel was exposed to a phosphoscreen for 1-2h at room temperature. The phosphoscreen was imaged on a Typhoon phosphorimager (Cytiva), and gel band intensity was quantified using ImageQuant TL (Cytiva).

### Western Blotting of EssR protein

*B. ovis* strains were grown on plain TSAB or TSAB supplemented with kanamycin for two days. Cells were collected and resuspended in sterile PBS to equivalent densities (measured optically at 600 nm). Cell resuspensions were mixed with an equal volume of SDS loading buffer and heated to 95°C for 5 minutes. 10 µl of each sample was loaded onto a 12.5% SDS-PAGE gel and resolved at 180V at room temperature. The proteins were transferred to a PVDF membrane (Millipore) using a semi-dry transfer apparatus at 10V for 30 minutes at room temperature. The membrane was blocked in 20 ml Blotto (Tris-buffered saline Tween-20 (TBST) + 5% milk) for 1h at room temperature. The membrane was blocked in 10 ml Blott + polyclonal rabbit α-EssR antiserum (1:1,000 dilution), and the membrane was incubated for 1h at room temperature. The membrane was washed three times with TBST. Goat α-rabbit IgG poly-horseradish peroxidase secondary antibody (Invitrogen; 1:10,000 dilution) was added to 10 ml Blotto and the membrane was incubated for 1h at room temperature. The membrane was washed three times with TBST, developed with ProSignal Pico ECL Spray (Prometheus Protein Biology Products) and imaged on a BioRad ChemiDoc Imaging System. Bands were quantified using ImageLab Software (BioRad).

### Western Blotting of CenR-3xFLAG protein

*B. ovis* strains carrying pQF plasmids were grown on TSAB supplemented with 30 µg/ml oxytetracycline for three days and samples for Western blot were collected and and prepared as described above. 10 µl of each sample was loaded onto a 12.5% SDS-PAGE containing 0.5% 2,2,2-Trichloroethanol (TCE) and resolved at 180V at room temperature. The gel was imaged on a BioRad Chemi-Doc Imaging system using the stain-free protein gel setting to activate TCE and visualize total proteins in the gel. The proteins were then transferred to a PVDF membrane (Millipore) using a semi-dry transfer apparatus at 10V for 30 minutes at room temperature and imaged under the stain-free blot setting on Chemi-Doc to visualize the total transferred proteins. The membrane was blocked in 20 ml Blotto (Tris-buffered saline Tween-20 (TBST) + 5% milk) for 1h at room temperature, then incubated in 15 ml Blotto + monoclonal α-FLAG antibody (Thermo; 1:10,000 dilution) for 1h at room temperature. The membrane was subsequently washed three times with TBST and goat α-mouse IgG poly-horseradish peroxidase secondary antibody (Thermo; 1:10,000 dilution) was added and incubated for 1h at room temperature. Membrane incubated with secondary antibody was subsequently washed three times with TBST, developed with ProSignal Pico ECL Spray (Prometheus Protein Biology Products) and imaged on ChemiDoc Imaging System. Bands were quantified using ImageLab Software (BioRad).

## Acknowledgements

We thank Maeve McLaughlin for experimental guidance on ChIP-seq protocol and analysis, and current and former Crosson Lab members for helpful discussions over the course of this study. We also thank Sundharraman Subramanian at the Cryo-EM Facility at Michigan State University for assistance with Cryo-EM. This study was supported by NIH award R35GM131762 to S.C.

## Supplemental Figures

**Figure S1.**
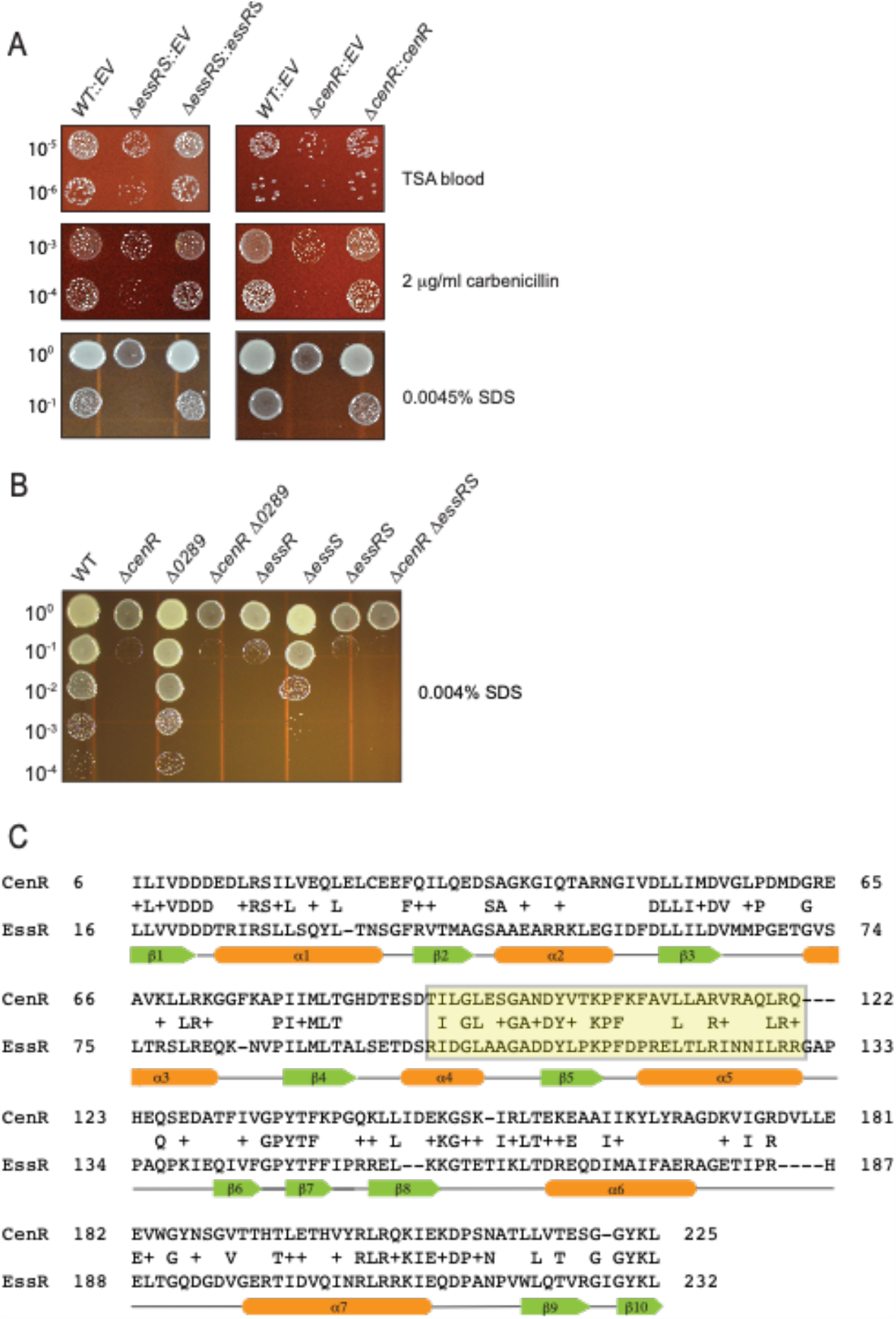
Genetic complementation of D*essRS* and D*cenR* mutants; SDS sensitivity assay at a reduced SDS concentration, and CenR-EssR sequence alignment. (A) Genetic complementation of *ΔessRS* and Δ*cenR* carbenicillin and SDS phenotypes. Complementing copies of the deleted genes were inserted into the ectopic *glmS* locus using Tn7. Dilution plating experiments were repeated at least three times for all strains. One representative experiment is shown. (B) SDS resistance phenotypes of strains harboring in-frame unmarked deletions (D) of *B. ovis* TCS gene loci *BOV_1929* (*cenR*), *BOV_0289, BOV_1472* (*essR*), *BOV_1473* (*essS*), alone and in combination show a larger dynamic range at an SDS concentration of 0.004% compared to 0.0045% (as presented in Figure 1). Dilution plating experiments were repeated at least three times for all strains, and one representative experiment is shown. (C) Amino acid sequence alignment of CenR and EssR. Amino acids highlighted in yellow show the primary structure of the α4-β5-α5 proteinprotein interaction interface predicted by AFComplex2 (see Figure 5C). Protein secondary structure is presented above the alignment (helix in orange; strand in green).

**Figure S2:**
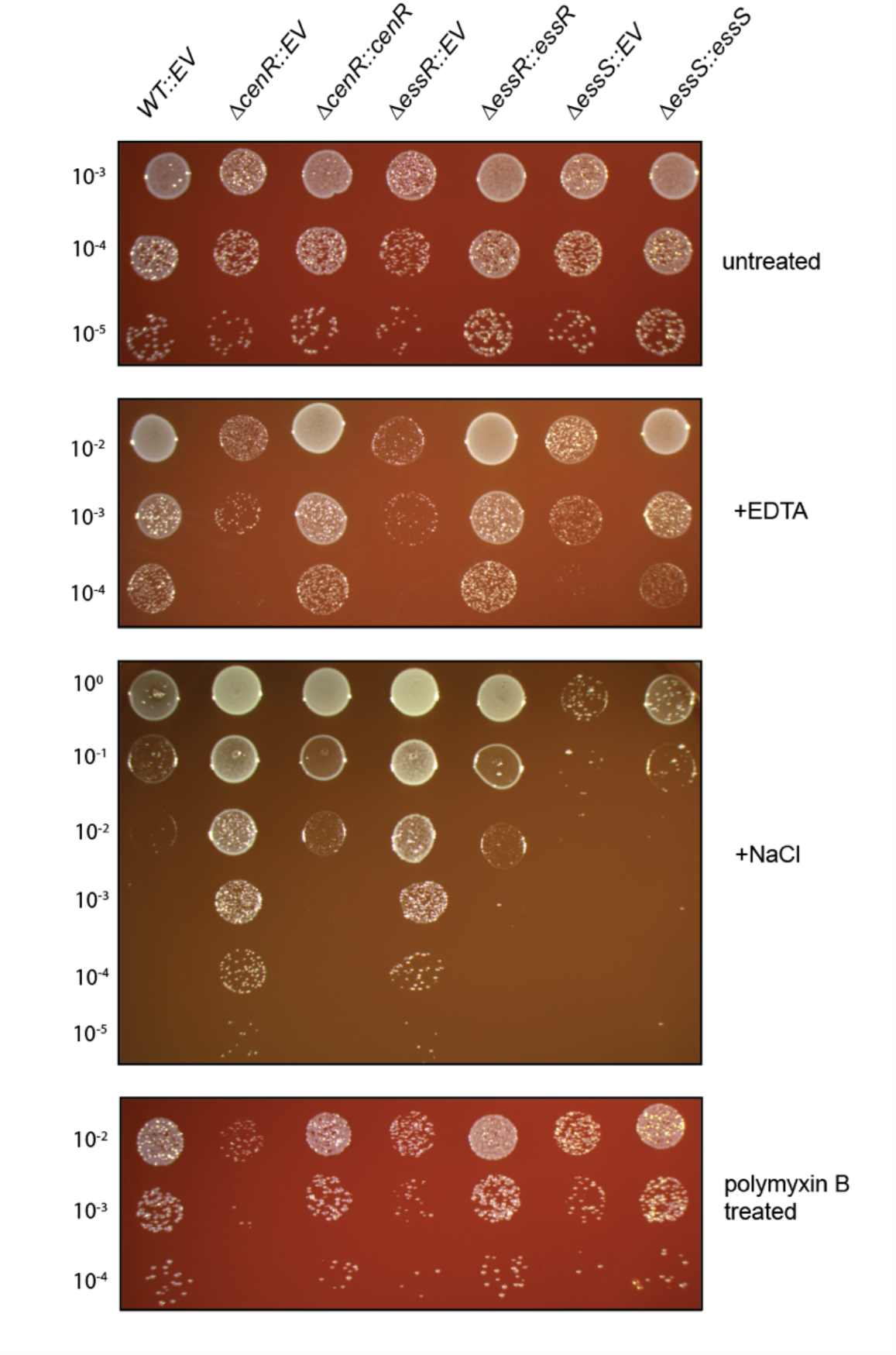
*cenR, essR*, and *essS* contribute to cell survival in the presence of diverse envelope stressors. Strains harboring in-frame unmarked deletions (D) of *cenR, essR, essS*, carrying integrated empty vectors (EV) or genetic complementation vectors (::*gene locus number*) were plated in log_10_ dilution series on plain TSA blood agar (untreated), TSA-B containing 2.75mM EDTA (+EDTA), TSA-B containing 215 mM NaCl (+NaCl), or were cells treated with 1 mg/mL polymyxin B for 80 minutes before being plated on TSA-B (polymyxin B treated). Dilution plating experiments were repeated three times for all strains, and one representative experiment is shown.

**Figure S3:**
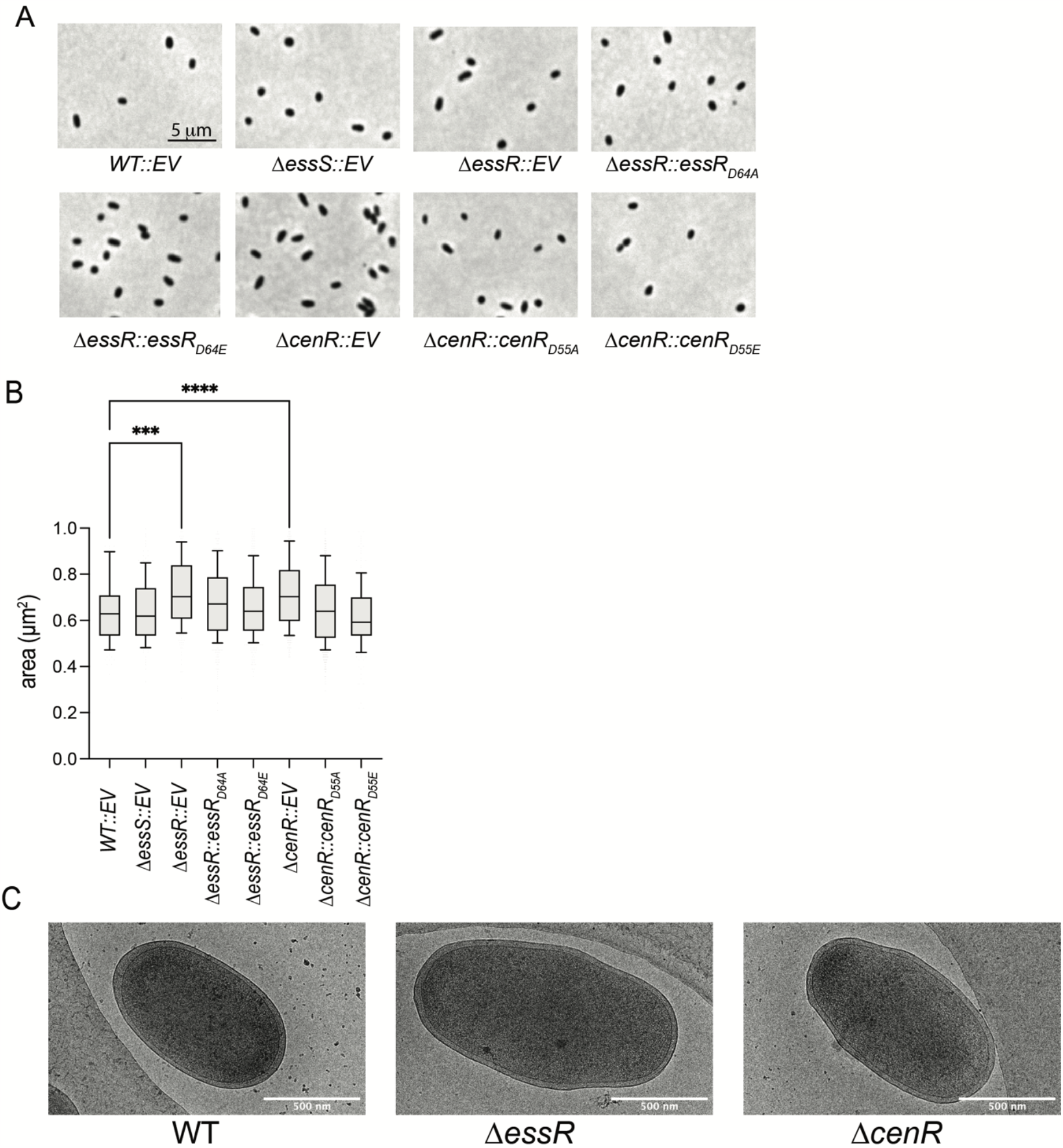
Deletion of *cenR* and *essR* results in increased cell area. A) Phase-contrast micrographs (630x magnification) of WT *B. ovis*, D*essS*, D*essR*, and D*cenR* in-frame deletion mutants; D*essR* complemented with *essR*_D64A_ and *essR*_D64E_; D*cenR* complemented with *cenR*_D55A_ and cen*R*_D55E_. B) Cell area analysis of WT (n=113), D*essS* (n=173), D*essR* (n=242), and D*cenR* (n=535) empty vector control strains (EV), D*essR* complemented with *essR*_D64A_ (n=478) and *essR*_D64E_ (n=466) alleles and D*cenR* complemented with *cenR*_D55A_ (n=908) and cen*R*_D55E_ (n=180) alleles. Mean is shown as a horizontal line in the box (25^th^-75^th^ percentile); whiskers capture from 10th-90^th^ percentile. Statistical significance was calculated using one-way ANOVA, followed by Dunnett’s multiple comparison test to WT empty vector control (WT::EV) (p < 0.001, ***; p < 0.0001, ****). C) Representative cryo-EM images of WT *B. ovis, ΔessR*, and *ΔcenR* in-frame deletion mutants.

**Figure S4:**
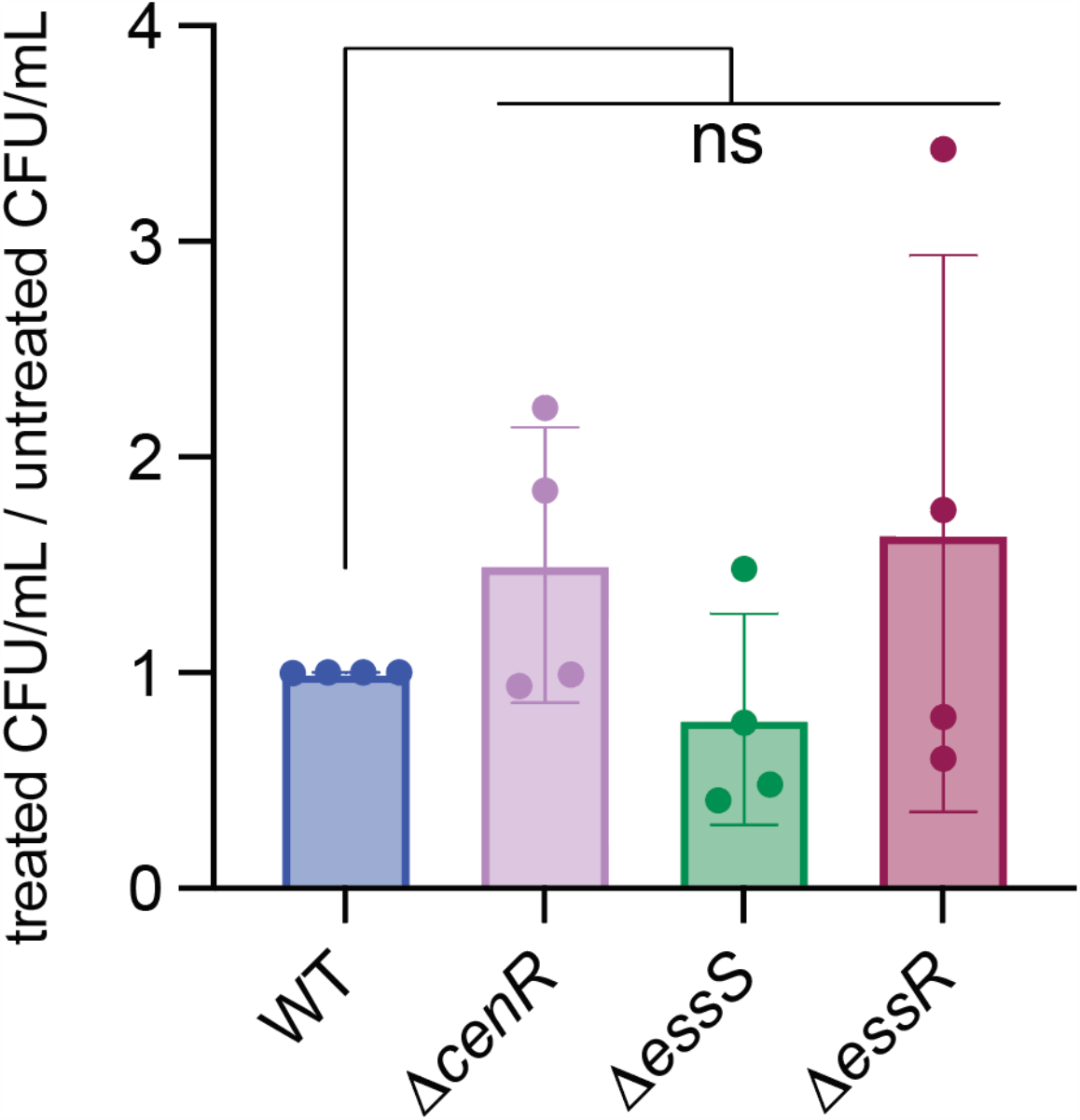
*ΔcenR, ΔessS*, and Δ*essR* are not sensitive to acidic pH. Strains harboring in-frame deletions of *cenR, essS*, and *essR* were incubated in Brucella broth at pH 7.0 or Brucella broth at pH 4.2 for 2 h before being serially diluted and plated on TSAB. CFU of treated cultures (pH 4.2) were normalized to CFU of untreated cultures (pH 7). Error bars represent the mean ± standard deviation of the four replicates. Statistical significance was calculated using one-way ANOVA, followed by Dunnett’s multiple comparisons test to WT (n.s. = non-significant, p > 0.05).

**Figure S5.**
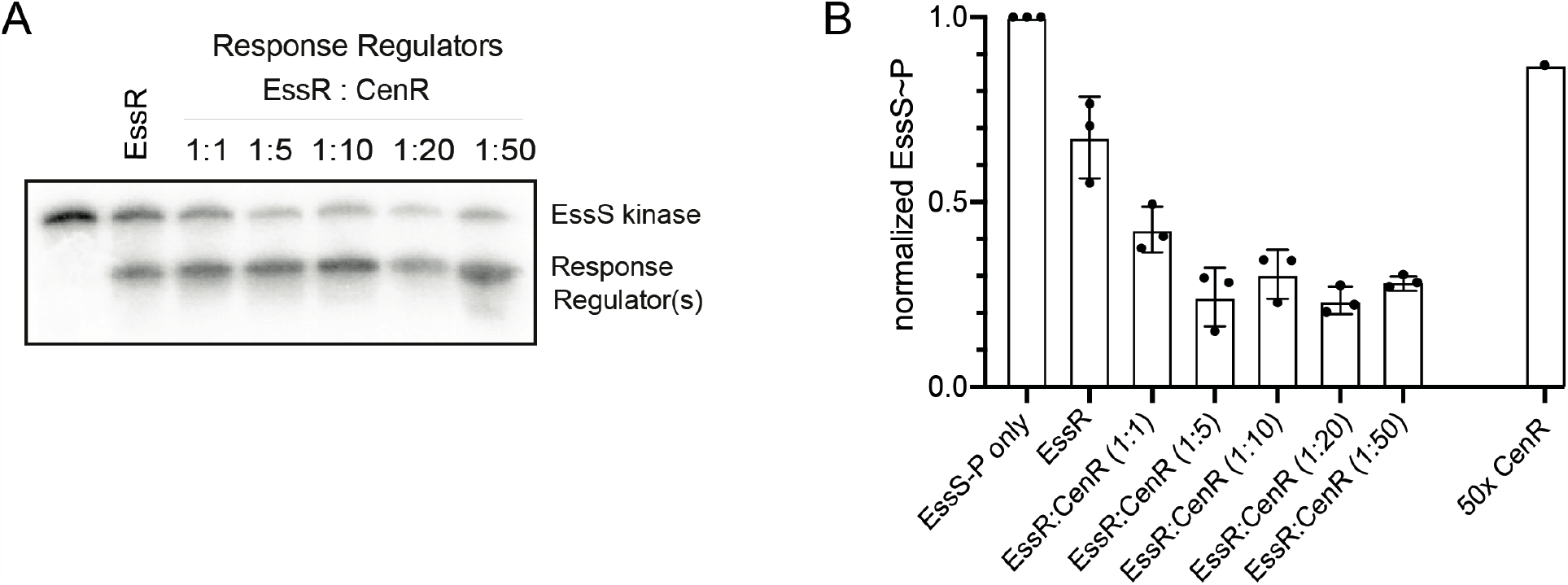
CenR enhances dephosphorylation of EssS∼P in the presence of EssR. A) *In vitro* phosphoryl transfer assay with purified EssS kinase domain and EssR (1:1) and increasing ratiometric amounts of CenR. All reactions were stopped after 20s. B) Quantification of EssS∼P levels; mean EssS∼P band intensity is set to 1 in the EssS∼P only reaction. Normalized EssS∼P levels 20s after addition of EssR and CenR in varying ratios is plotted. Error bars represent the standard deviation of three biological replicates.

**Figure S6:**
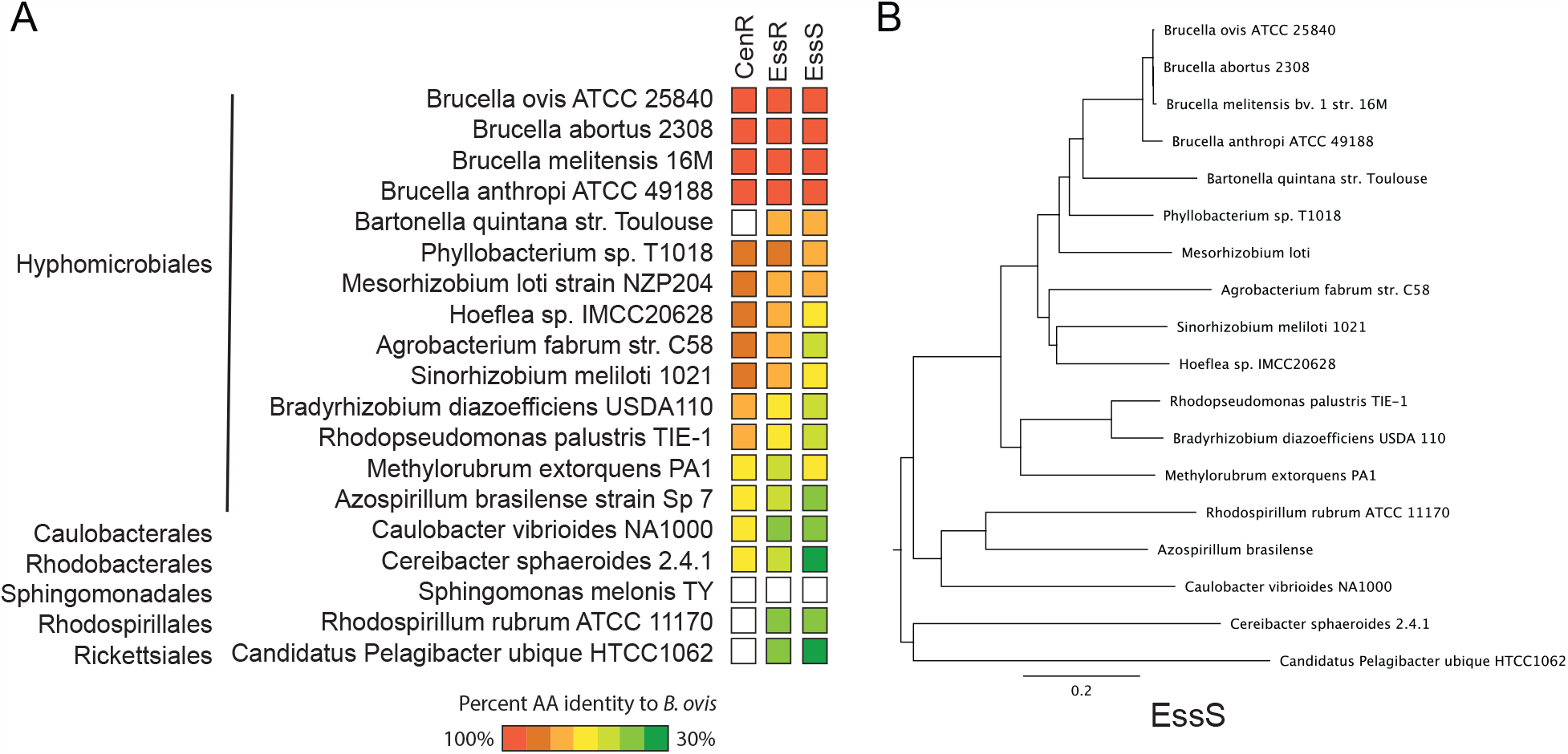
CenR, EssR, and EssS are widely distributed in the class Alphaproteobacteria. A) Representative Alphaproteobacterial genomes were queried for reciprocal top BLAST hits of CenR, EssR, and EssS. A gene was determined as present (colored box) if a reciprocal top-hit pair was found, and as absent (white) if no such pair was found. The color of the boxes reflects the percent amino acid (AA) identity (over the full protein length) to the corresponding *B. ovis* sequences. B) A phylogenetic tree based on the amino acid sequence of EssS orthologs. Global alignment with free end gaps was used as the alignment type and Blosum62 as the cost matrix. Jukes-Cantor was used as the genetic distance model and neighbor-joining was the tree build method with no outgroup.

