## Supplementary material for "Cross regulation in a three-component cell envelope stress signaling system of *Brucella*": Table S2

Table S2. Two-component system (TCS) genes in *Brucella* *ovis* ATCC 25840

| **BOV locus** | **NCBI RefSeq locus** | **Gene name(s)** | **Domains** | **Essential^1^** | **Mutants generated?** |
| --- | --- | --- | --- | --- | --- |
| BOV_0099 | BOV_RS00485 | - | REC | N | Y |
| BOV_0128 | BOV_RS00640 | *regB/prrB* | TM-HK | N | N |
| BOV_0131 | BOV_RS00655 | *regA*/*prrA* | REC-DBD | N | Y |
| BOV_0190 | BOV_RS18300/  BOV_RS00940^2^ | - | Pseudo REC-DBD | N | N |
| BOV_0289 | BOV_RS01475 | - | TM-HK | N | Y |
| BOV_0312 | BOV_RS01585 | *phoR* | TM-HK | N | Y |
| BOV_0331 | BOV_RS01685 | *prlR* | REC-DBD | N | Y |
| BOV_0332 | BOV_RS16185^2^ | *prlS* | Pseudo hybrid HK-REC | N | Y |
| BOV_0357 | BOV_RS01805 | - | TM-Cache-HK | N | Y |
| BOV_0358 | BOV_RS01810 | - | REC-DBD | N | Y |
| BOV_0577 | BOV_RS02880 | *divJ* | TM-HK | N | Y |
| BOV_0603 | BOV_RS03015 | *feuP* | REC-DBD | N | Y |
| BOV_0604 | BOV_RS03020 | *feuQ* | TM-HK | N | Y |
| BOV_0611 | BOV_RS03055 | - | REC-DBD | N | Y |
| BOV_0612 | BOV_RS03060 | - | TM-HK | N | Y |
| BOV_0615 | BOV_RS03075 | *pleC* | TM-PAS-HK | N | Y |
| BOV_1004 | BOV_RS04980 | *cckA* | TM-PAS-HK-REC | Y | N |
| BOV_1073 | BOV_RS05335 | *ntrX* | REC-AAA+ ATPase-DBD | Y | N |
| BOV_1074 | BOV_RS05340 | *ntrY* | TM-PAS-HK | N | N |
| BOV_1075 | BOV_RS05345 | *ntrC* | REC-AAA+ ATPase-DBD | N | Y |
| BOV_1076 | BOV_RS05350 | *ntrB* | HK | N | Y |
| BOV_1472 | BOV_RS07250 | *essR* | REC- DBD | N | Y |
| BOV_1473 | BOV_RS07255 | *essS* | TM-HK | N | Y |
| BOV_1541 | BOV_RS07590 | *chpT* | HPT | Y | N |
| BOV_1542 | BOV_RS07595 | *ctrA* | REC-DBD | Y | N |
| BOV_1549 | BOV_RS07630 | *pdhS* | PAS-HK | Y | N |
| BOV_1602 | BOV_RS07885 | - | HWE-HK | N | Y |
| BOV_1604 | BOV_RS07895 | *phyR* | REC | N | Y |
| BOV_1607 | BOV_RS07910 | *phyT* | TM-CHASE3-HWE-HK | N | N |
| BOV_1929 | BOV_RS09460 | *cenR*/*otpR* | REC-DBD | N | Y |
| BOV_2010 | BOV_RS09885 | *bvrR* | REC-DBD | Y | N |
| BOV_2011 | BOV_RS09890 | *bvrS* | TM-HK | Y | N |
| BOV_2016 | BOV_RS09915 | *divL* | PAS-HK | Y | N |
| BOV_2058 | BOV_RS10125 | *phoB* | REC-DBD | N | Y |
| BOV_A0037 | BOV_RS10530 | *nodV* | TM-HK | N | Y |
| BOV_A0038 | BOV_RS10535 | *nodW* | REC-DBD | N | Y |
| BOV_A0040 | BOV_RS10545 | *cpdR* | REC | N | N |
| BOV_A0077 | BOV_RS10730^2^ | - | Pseudo REC-AAA+ ATPase-DBD | N | N |
| BOV_A0209 | BOV_RS11400 | - | TM-HK | N | Y |
| BOV_A0210 | BOV_RS11405 | - | REC-DBD | N | Y |
| BOV_A0358 | BOV_RS12165 | - | REC-DBD | N | Y |
| BOV_A0412 | BOV_RS12435^2^ | - | Pseudo HK | N | Y |
| BOV_A0413 | BOV_RS12440 | - | REC-DBD | N | Y |
| BOV_A0554 | BOV_RS13160 | *lovhK* | PAS-HWE-HK | N | Y |
| BOV_A0575 | BOV_RS13265 | *pleD* | REC-GGDEF | N | Y |
| BOV_A0576 | BOV_RS13270 | *divK* | REC | Y | N |
| BOV_A1045 | BOV_RS15645 | *ftcR* | REC-DBD | N | Y |

Abbreviation keys:

TM: transmembrane	HWE-HK or HK: histidine kinase

REC: response regulator receiver domain	DBD: DNA-binding domain

AAA+ ATPase: sigma 54 activated domain	GGDEF: diguanylate cyclase domain

PAS/CHASE3/CACHE: sensory domains	HPT: histidine phosphotransferase

^1^Essential genes based on Gumbel and HMM essential gene calculations in TRANSIT [1], using the Tn-Himar sequencing data available at accession SRR19632676 in the NCBI Sequence Read Archive. Genes called essential using both the Gumbel and HMM approaches are noted.

^2^Annotated as pseudogene in RefSeq based on the presence of insertion/deletion or nonsense mutations relative to other *Brucella* species.

1.	DeJesus, M.A., et al., *TRANSIT--A Software Tool for Himar1 TnSeq Analysis.* PLoS Comput Biol, 2015. **11**(10): p. e1004401.
